# Biofunctional matrix models reveal mineral-dependent mechanoregulation of bone metastatic breast cancer

**DOI:** 10.1101/2022.06.29.498001

**Authors:** Siyoung Choi, Matthew A. Whitman, Adrian A. Shimpi, Nicole D. Sempertegui, Aaron Chiou, Joseph E. Druso, Akanksha Verma, Stephanie C. Lux, Zhu Cheng, Matthew Paszek, Olivier Elemento, Lara A. Estroff, Claudia Fischbach

## Abstract

Bone metastasis is a leading cause of breast cancer-related deaths and often initiated by tumor cell dissemination to osteogenic niches. During new bone formation, osteoblasts first deposit osteoid, the collagen I-rich, unmineralized component of bone ECM, within which carbonated hydroxyapatite nanoparticles subsequently form. However, it remains elusive how bone matrix mineralization dictates tumor cell phenotype due in part to the lack of relevant model systems. Using biofunctional, collagen I-based bone matrix models with physiological, intrafibrillar mineralization, we show that mineralization inhibits proliferation, while inducing a stem-like phenotype in tumor cells. These changes were due to reduced mechanosignaling contradicting the conventional assumption that increased rigidity caused by mineralization stimulates metastatic progression. Our findings are translationally relevant as the presence of mineral reduced tumor growth *in vivo* and upregulated a gene signature that correlated with decreased patient mortality. Our results could help explain why decreased bone mineral density increases the risk for bone metastasis in patients and highlight that bone metastasis models should integrate organic and inorganic matrix components in a manner that mimics physiological mineralization.

## Introduction

Breast cancer frequently metastasizes to bone where it leads to osteolysis and poor clinical prognosis^1–3^, but therapies to treat or prevent bone metastasis are lacking. Most prior research has focused on clinically apparent bone degradation uncovering that unbalanced osteoclast activity and consequential activation of the ‘vicious cycle of bone metastasis’ allows asymptomatic, residual disease to develop into overt metastasis^4, 5^. However, how tumor cells initially colonize and survive in the skeletal microenvironment is poorly understood. Indeed, tumor cells can disseminate to bone very early during tumor development and enter a state of latency in which they proliferate less and assume stem cell-like properties^6–8^. Activation of these latent tumor cells to a more proliferative phenotype contributes to their outgrowth into metastatic tumors after years or even decades. As microenvironmental conditions are key determinants of tumor cell latency and stemness^7, 9^, it will be critical to better define how bone-specific niches and changes thereof influence the stem-like phenotype of tumor cells in the skeleton.

While most research on early-stage bone metastasis focuses on the marrow, tumor cells also colonize osteogenic niches in the skeleton. These niches consist of osteoblasts and their progenitors whose hallmark function is to deposit and mineralize extracellular matrix (ECM)^9, 10^ (Fig. 1a). Conditions affecting osteoblasts and their progenitors (e.g., aging or chemotherapy) inhibit the formation of mineralized matrix and alter the secretome of osteoblast-lineage cells^11, 12^. While the latter has been shown to regulate metastatic latency^13^ and outgrowth^11, 12^, it remains underappreciated how varied bone ECM formation and mineralization influence tumor cell phenotype. Better understanding these connections will be important as changes of the ECM play a key role in regulating tumor cell phenotype, because disseminated cancer cells directly interact with the matrix^10, 14^, and because breast cancer patients with decreased bone mineral density have a higher risk of developing bone metastasis^15, 16^.

**Figure 1:**
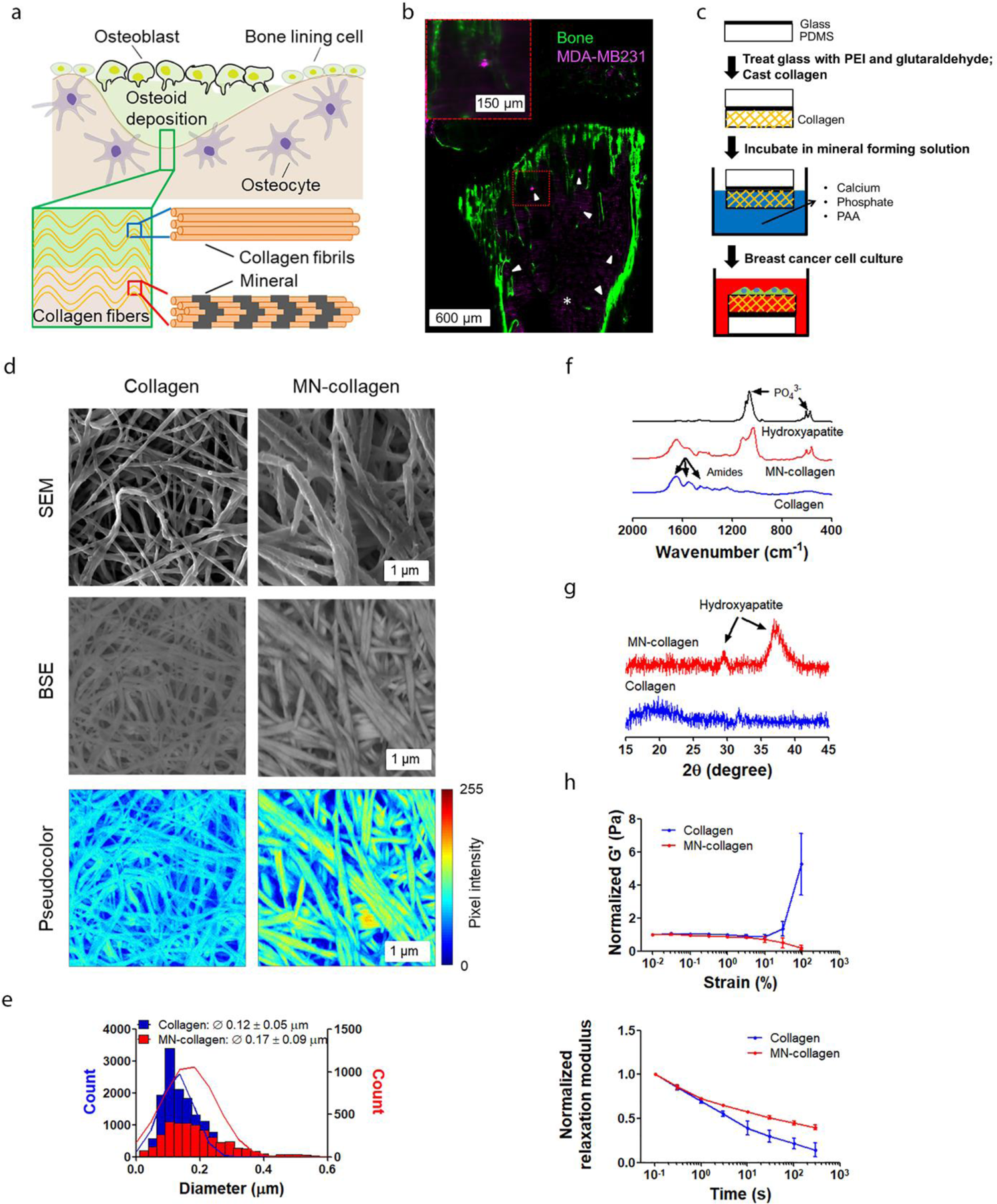
Engineered bone matrix models to study tumor cell interactions with mineralized collagen. **a,** Schematic of bone ECM formation in osteogenic niches. Osteoblasts first deposit osteoid, which consists primarily of collagen type I fibers (blue box) that become mineralized (red box) over time by intrafibrillar mineralization. **b,** Intracardially injected MDA-MB231 labeled with Cy5-containing silica nanoparticles (magenta) disseminate to regions in which new bone matrix becomes mineralized as visualized by light sheet imaging of a calcein-labeled, cleared mouse tibia (green). White arrow-heads and * indicate MDA-MB231 cells and autofluorescence of bone marrow, respectively. Scale bar = 600 µm and 150 µm (inset). **c**, Experimental setup to mineralize fibrillar collagen type I matrices via the PILP method for subsequent analysis of breast cancer cell phenotype. PDMS: Polydimethylsiloxane; PEI: polyethyleneimine; PAA: polyaspartic acid. **d**, Representative SEM and BSE images visualizing the fibrillar nature and mineral content of collagen and mineralized collagen (MN-collagen) substrates. Pseudocolor indicates mineral content as determined from BSE images. Scale bar = 1 µm. **e**, Fibril diameter (Ø) of control and mineralized collagen substrates quantified from SEM images. **f**, FT-IR spectra of the different collagen substrates and highly crystalline hydroxyapatite particles as control. Arrows indicate phosphate (450 - 750 cm^-1^ and 900 - 1300 cm^-1^) and amide peak areas (1200 - 1700 cm^-1^). **g**, pXRD patterns indicate the presence of hydroxyapatite in mineralized, but not control collagen. **h**, Mechanical properties of collagen and mineralized collagen. Strain sweeps with normalized storage modulus (G’) for strain stiffening measurement (top) and time sweeps with normalized relaxation modulus for stress relaxation measurement (bottom) (*n* = 3).

During osteogenesis, osteoblasts first deposit osteoid, the collagen I-rich, unmineralized component of bone ECM, within which carbonated hydroxyapatite (HA; Ca_10_(PO_4_CO_3_)_6_(OH)_2_) nanoparticles subsequently form^17, 18^ (Fig. 1a). This process of mineralization is accompanied by distinct changes in bone matrix mechanical properties (e.g., increased stiffness, reduced stress relaxation^19^), which can impact tumor cell phenotype through altered mechanosignaling^20–22^. Moreover, bone matrix mineralization is a dynamic process that changes as a function of anatomical site, age, diet, and disease^23–26^. How changes in bone matrix mineralization influence the phenotype of tumor cells, however, remains unclear due in part to a lack of model systems that allow selective control over bone matrix mineral content for mechanistic studies. Studies with such systems will be critical to elucidate why reduced bone formation and local mineral density increase the risk for bone metastasis^12, 27^, whereas conditions promoting new bone formation and mineralization inhibit bone metastasis^28, 29^.

Conventional models of breast cancer (i.e., 2-D cell culture and mouse models) fail to recapitulate *(i)* compositional, structural, and mechanical alterations of bone matrix and *(ii)* do not allow control over the level of mineralization that changes during osteogenesis and in response to conditions that correlate with increased risk for bone metastasis (e.g. age, Vit. D deficiency, premetastatic bone remodeling)^24, 25, 30, 31^. More advanced methods such as culturing fragments of tumor cell-containing mouse bones *ex vivo* can recapitulate tumor cell interactions with bone matrix in the presence of bone-resident tumor cells, but matrix mineralization cannot be controlled independently in this system^32^. While many protocols have been developed to mineralize biomaterials for cell culture studies or regenerative applications, most of these methods deposit HA on top of substrates rather than mimicking physiological mineralization. Yet, bone matrix is a composite material whose unique mechanical properties are determined by HA-enforced collagen fibers that form by intrafibrillar nucleation of the mineral phase within the organic matrix^19, 33^.

Here, we have developed osteoid-like and bone-like scaffolds in which we can selectively adjust bone matrix mineral content for both *in vitro* and *in vivo* experiments in a physiologically relevant manner. These model systems include: *(i)* 2.5-D collagen matrices in which we synthetically induce bone-like intrafibrillar mineralization through a modified polymer-induced liquid precursor (PILP) method^34^ and *(ii)* decellularized trabecular bone scaffolds in which we can selectively remove mineral without affecting organic ECM composition and microstructure. Using these biofunctional scaffolds, we probed the role of bone matrix mineralization in regulating the phenotype of breast cancer cells *in vitro* and tested which role mechanosignaling plays in this process. Then, we evaluated the impact of our results on tumor growth in mice and on patient prognosis using bioinformatic analysis of published clinical data sets. Collectively, our results suggest that the mineral content of bone matrix regulates skeletal metastasis by influencing the phenotype of breast cancer cells. These findings underscore that bone matrix is an important component of the osteogenic niche that can impact bone metastasis and thus, should be considered when designing model systems for studies of breast cancer bone metastasis.

## Results

### Disseminated tumor cells seed mineralizing bone niches, which can be mimicked by intrafibrillar mineralization of collagen

Whether disseminated tumor cells localize to skeletal regions of new matrix mineralization is poorly characterized given the intrinsic difficulty of imaging rare numbers of tumor cells within fully intact, highly autofluorescent, mineralized bones. To circumvent this challenge, we have labeled MDA-MB231 breast cancer cells with bright fluorescent silica nanoparticles and intracardially injected them into athymic nude mice in which newly mineralizing bone surfaces were labeled with calcein^35^. As expected, cancer cells localized to niches within the marrow in this model^36^. However, light sheet microscopy of cleared mouse tibiae also revealed that tumor cells co-localized with regions of new matrix mineralization (Fig. 1b) consistent with previous results identifying the osteogenic niche as a target for early-stage bone colonization^10^.

To investigate whether bone matrix mineralization alone affects the phenotype of tumor cells, we utilized a bone-like ECM platform in which we induced biofunctional intrafibrillar mineralization of collagen using a modified polymer-induced liquid-precursor (PILP) method^34, 37^. We chose collagen type I for this platform as osteoid primarily consists of collagen type I and because bone matrix mineralization primarily occurs in collagen type I fibers^38^. Briefly, collagen was cast into polydimethylsiloxane (PDMS) microwells or on glass coverslips and mineralized in a solution containing calcium and phosphate ions as well as polyaspartic acid to control the formation of HA nanocrystals within collagen fibrils (Fig. 1c). In this system, the polyaspartic acid serves a function similar to non-collagenous proteins in bone, resulting in the formation of bone-like mineralized collagen fibers^20^. Scanning electron microscopy (SEM) image analysis confirmed that mineralization did not impact the fibrous architecture of collagen but increased the thickness of individual fibrils (Fig. 1d, e) due to mineral formation as confirmed by backscattered electron (BSE) imaging (Fig. 1d). Fourier transform infrared (FT-IR) spectroscopy and powder X-ray diffraction (pXRD) further identified that the chemical composition and phase of the newly formed mineral were consistent with HA (Fig. 1f, g). FT-IR spectra of both control and mineralized collagen contained characteristic protein amide peaks between 1200 - 1700 cm^-1^, while PO_4_^3-^ peaks indicative of HA (500-700 cm^-1^ and 900-1200 cm^-1^) were only detected in mineralized collagen (Fig. 1f). Moreover, the mineral content of these matrices was comparable to human bone as suggested by FT-IR analysis of the mineral to matrix ratio (Supplementary Fig. 1; mineralized collagen: approx. 4; human bone: 3-6)^26^. The phase of the mineral that formed in collagen was similar to the poorly crystalline, non-stoichiometric carbonated apatite in bone^39^ as indicated by the presence and broadening of pXRD peaks at 26° and 32° (Fig 1g). Collagen is a viscoelastic material that undergoes non-linear elastic changes upon deformation. Upon mineralization, the non-linear elastic response of the collagen will change, and thus cellular behavior is likely to change also^40^. Indeed, mineralization increases the storage and loss moduli of collagen indicating increased stiffness as expected (Supplementary Fig. 2). In addition, rheological analysis revealed that mineralization prevents collagen strain stiffening while reducing collagen stress relaxation in response to strain (Fig. 1h). Taken together, these results indicate that bone-like collagen-HA composites can be formed by synthetic intrafibrillar collagen mineralization and that the resulting substrates are characterized by altered chemical, physical, and mechanical properties.

### Collagen mineralization alters breast cancer cell gene expression and growth *in vitro*

As microfabricated collagen and mineralized collagen substrates recapitulated the two extremes of bone matrix mineralization (i.e., non-mineralized osteoid vs. mineralized bone matrix), we next used these materials to test how bone matrix mineralization affects tumor cell phenotype. When cultured on mineralized versus control collagen metastatic MDA-MB231 breast cancer cells spread less, were rounder, and exhibited reduced remodeling of their surrounding collagen network as indicated by limited fiber alignment and densification that is known to mediate local strain stiffening in control collagen^41^ (Fig. 2a, b). Consistent with these morphological differences principal component analysis (PCA) of bulk RNA-seq data sets suggested that collagen mineralization globally affects gene expression of MDA-MB231 (Fig. 2c). Differential gene expression analysis further identified that genes associated with breast cancer metastasis (ANGPTL2, CAMK1D, CEMIP)^42–44^ and stemness (B4GALNT3, TRIB3)^45, 46^ were among the top 50 differentially expressed genes in the mineralized collagen condition, while most of the top 50 ranked genes regulated by collagen were linked with cell cycle (CDCA7, CHTF18, DBF4B, KIF18B) (Fig. 2d, Table 1, Supplementary Fig. 3). Gene set enrichment analysis (GSEA) confirmed that cell cycle-related gene sets, most prominently DNA replication, were over-represented in cells cultured on collagen, while gene sets associated with stress response and ECM disassembly, transcription, and cell migration were over-represented in cells cultured on mineralized collagen (Fig. 2e, Supplementary Fig. 4). These results suggest that bone matrix mineralization impacts breast cancer cell functions centrally implicated in bone metastasis.

**Figure 2:**
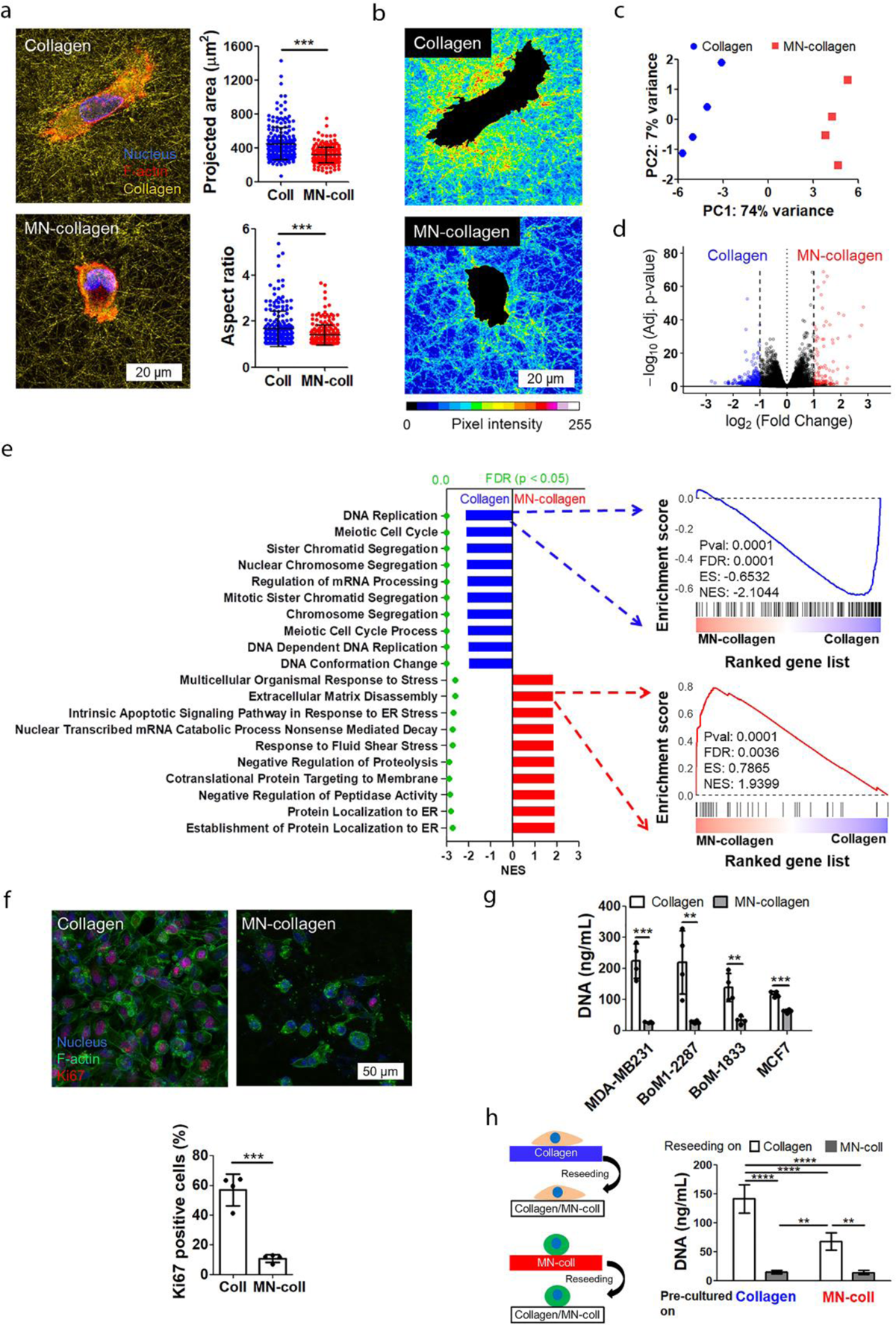
Mineralization of collagen alters breast cancer cell gene expression and growth. **a,** MDA-MB231 morphology and collagen network structure 5 h after seeding visualized via confocal reflectance microscopy (left) and analysis of projected cell area and aspect ratio (right). **b,** Pseudo-colored reflectance intensity maps indicate differences in collagen remodeling between both conditions. Scale bar = 20 µm. **c-e,** Effect of collagen mineralization on MDA-MB231 gene expression determined by principal component analysis (PCA) (c), differential gene expression analysis (*n* = 4) (d), and gene set enrichment analysis (GSEA) (e) of bulk RNA-seq data collected after 7 d of culture on the different substrates. In d, genes colored in red are upregulated by cells cultured on mineralized collagen, while genes colored in blue are upregulated on collagen. For GSEA, the top 10 enriched pathways in cells cultured on collagen and mineralized collagen were derived from GO biological process. Green dots indicate false discovery rates (FDR). Normalized enrichment scores (ES) are denoted by NES. **f**, Representative confocal micrographs and corresponding quantification of Ki67^+^ MDA-MB231 cells. **g,** fluorimetric quantification of DNA content of breast cancer cell lines cultured for 7 d on the different substrates (*n* = 4). **h**, Analysis of MDA-MB231 growth following 7 days of pre-culture on the different substrates (*n* = 4). **: *P* < 0.01, ***: *P* < 0.001, ****: *P* < 0.0001.

**Table 1.** Top 50 differentially expressed genes between collagen and mineralized collagen (p-value < 0.05)

| Top 50 Ranked Genes |  |  |  |
| --- | --- | --- | --- |
| Mineralized Collagen |  | Collagen |  |
| ABCA3 | NEDD9 | AC006058.1 | KIF23 |
| ADM2 | NNMT | AC008735.2 | L3HYPDH |
| ANGPTL2 | NUPR1 | AC068831.7 | LAPTM5 |
| AQP1 | PDLIM4 | AHSA2P | LUC7L3 |
| ATF3 | PLAT | AJUBA | MAMDC4 |
| B4GALNT3 | PRSS2 | AL136164.4 | MAP7D3 |
| BHLHE40 | PRSS3 | ALS2CL | MIR222HG |
| BMF | PTGS2 | ARGLU1 | MIR503HG |
| CAMK1D | RRAD | ARHGAP33 | NSUN5P1 |
| CEMIP | SELENOM | CARS2 | PABPC1L |
| CHAC1 | SGK1 | CASP2 | PLEKHA7 |
| CRIP2 | SLC43A2 | CCDC14 | PLK4 |
| CTSF | SLFN5 | CDCA7 | PNISR |
| EPHB6 | SMIM14 | CDH24 | PNN |
| FBN2 | SQSTM1 | CDK5RAP3 | RAD54L |
| FOS | TAGLN2 | CHTF18 | RECQL4 |
| FOSB | TCP11L2 | CREBZF | RSRP1 |
| HDAC5 | TMEM45B | DBF4B | SEMA7A |
| IGFBP1 | TPPP | DDX12P | SERINC2 |
| KLHL24 | TPPP3 | DDX39B | SLC25A27 |
| KRT19 | TRIB3 | FAM193B | SUGP2 |
| LGALS3 | TSC22D1 | HJURP | TGM2 |
| MEF2C | TSC22D3 | IL11 | TROAP |
| MRPL41 | TYMP | KIF18B | VAMP1 |
| NBL1 | VWA1 | KIF20A | ZNF692 |

As gene sets related to cell-cycle related pathways were most prominently affected by changes in collagen mineralization, we next measured tumor cell growth on the different substrates. Both MDA-MB231 Ki67 positivity and bromodeoxyuridine (BrdU) incorporation as well as DNA content were reduced by culture on mineralized collagen relative to collagen, indicating that collagen mineralization decreases tumor cell proliferation (Fig. 2f, g, Supplementary Fig. 5). Accordingly, collagen mineralization reduced growth of the bone tropic MDA-MB231 subclones BoM1-2287 and BoM-1833^20^, and estrogen-receptor positive MCF-7 cells supporting that our findings were not limited to MDA-MB231 (Fig. 2g). Patients with metastatic breast cancer have high plasma fibronectin concentrations (300 µg/mL) and fibronectin readily adsorbs onto the bone mineral hydroxyapatite^47, 48^. However, differences in fibronectin adsorption did not contribute to our results as validated in a control experiment in which we preadsorbed relevant fibronectin concentrations to the different substrates (Supplementary Fig. 6). Next, we evaluated if this decreased proliferative state is reversible in scenarios that lead to decreased matrix mineralization (e.g. aging, chemotherapy, osteolysis)^24, 25, 49^. To this end, we precultured MDA-MB231 on collagen and mineralized collagen for 7 days, a timeframe reported to cause irreversible phenotypic changes^50, 51^, and then reseeded them onto either collagen or mineralized collagen (Fig. 2h). Indeed, cells pre-cultured on mineralized collagen assumed a more proliferative phenotype when reseeded onto non-mineralized collagen albeit at reduced levels relative to cells that were precultured on collagen. In contrast, reseeding cells onto mineralized collagen significantly decreased tumor cell growth irrespective of the substrate on which cells were cultured previously (Fig. 2h). These data indicate that collagen mineralization reduces tumor cell growth in a reversible manner suggesting that quiescent cells may develop into proliferative lesions as matrix mineralization changes.

In osteogenic niches *in vivo*, tumor cells not only interact with matrix, but also with osteoblasts and their progenitors. Therefore, we next validated that increased matrix mineralization correlates with decreased tumor cell growth in the presence of bone-resident cells. Indeed, MDA-MB231 co-cultured with human bone marrow-derived mesenchymal cells (hMSCs) that had deposited mineralized matrix grew less than tumor cells co-cultured with hMSCs that failed to mineralize matrix (Supplementary Fig. 7c). While varied mineralization in these co-culture experiments was mimicked by the presence or absence of osteogenic media, similar scenarios may occur *in vivo* because tumor cells secrete factors that alter bone growth^52, 53^ and inhibit the formation of mineralized bone matrix by hMSCs (Supplementary Fig. 7a). Importantly, in scenarios where mineral matrix formation by hMSCs was inhibited (either by lack of osteogenic factors or the presence of tumor-secreted factors), hMSC deposited increased amounts of fibronectin (Supplementary Fig. 7b), a matrix protein that can independently promote bone metastasis and may thus exaggerate differences in scenarios in which bone matrix formation is inhibited *in vivo*. Collectively, these results suggest that changes in bone matrix mineralization correlate with altered tumor cell growth and are relevant or even amplified in the presence of bone-resident cells.

### Mineralized collagen increases a stem-like phenotype in breast cancer cells

Motivated by our observation that collagen mineralization reduces tumor cell proliferation (Fig. 2), a hallmark of quiescent stem-like tumor cells^7, 54^, and because dissemination to bone confers stem-like properties on cancer cells^55^, we next determined whether mineralized collagen regulates the stem-like phenotype of breast cancer cells. These experiments were performed with an MDA-MB231 stem cell reporter cell line that expresses GFP as a function of Nanog promoter activity^56^, which we validated to correlate with expression of two other stem cell markers, Oct4 and Sox2 (Fig. 3a). Flow cytometry and confocal image analysis indicated that culture on mineralized collagen increased the number of GFP^high^ cells relative to culture on collagen or polystyrene (PS) (Fig. 3a, b). To confirm these changes were driven by mineral, we performed a dose response study in which substrates of controlled HA content were fabricated by dissolving HA from fully mineralized collagen at physiological pH for up to 6 days, a process that selectively removed HA without affecting collagen microstructure (Fig. 3c, Supplementary Fig. 8). Decreasing mineral content reduced the percentage of Nanog-GFP positive cells supporting that collagen mineralization directly regulates the stem-like phenotype of breast cancer cells (Fig. 3d). Further validating the induction of stemness by mineral, parental MDA-MB231 cultured on mineralized collagen also exhibited increased activity of the stem cell marker aldehyde dehydrogenase (ALDH)^57, 58^ (Fig. 3e). Moreover, MDA-MB231 precultured on mineralized collagen formed more colonies in soft agar relative to the same cells precultured on PS or collagen indicating that interactions with mineralized collagen endow tumor cells with survival advantages in adhesion-independent scenarios (Fig. 3f). Bioinformatic analysis of our RNA-seq data set suggested that upregulation of *FOXD3* target genes may be involved, a transcription factors that regulates stem cell pluripotency and self-renewal^59, 60^, reduces proliferation of breast cancer cells, and predicts better survival^61^ (Supplementary Fig. 4a, b).

**Figure 3:**
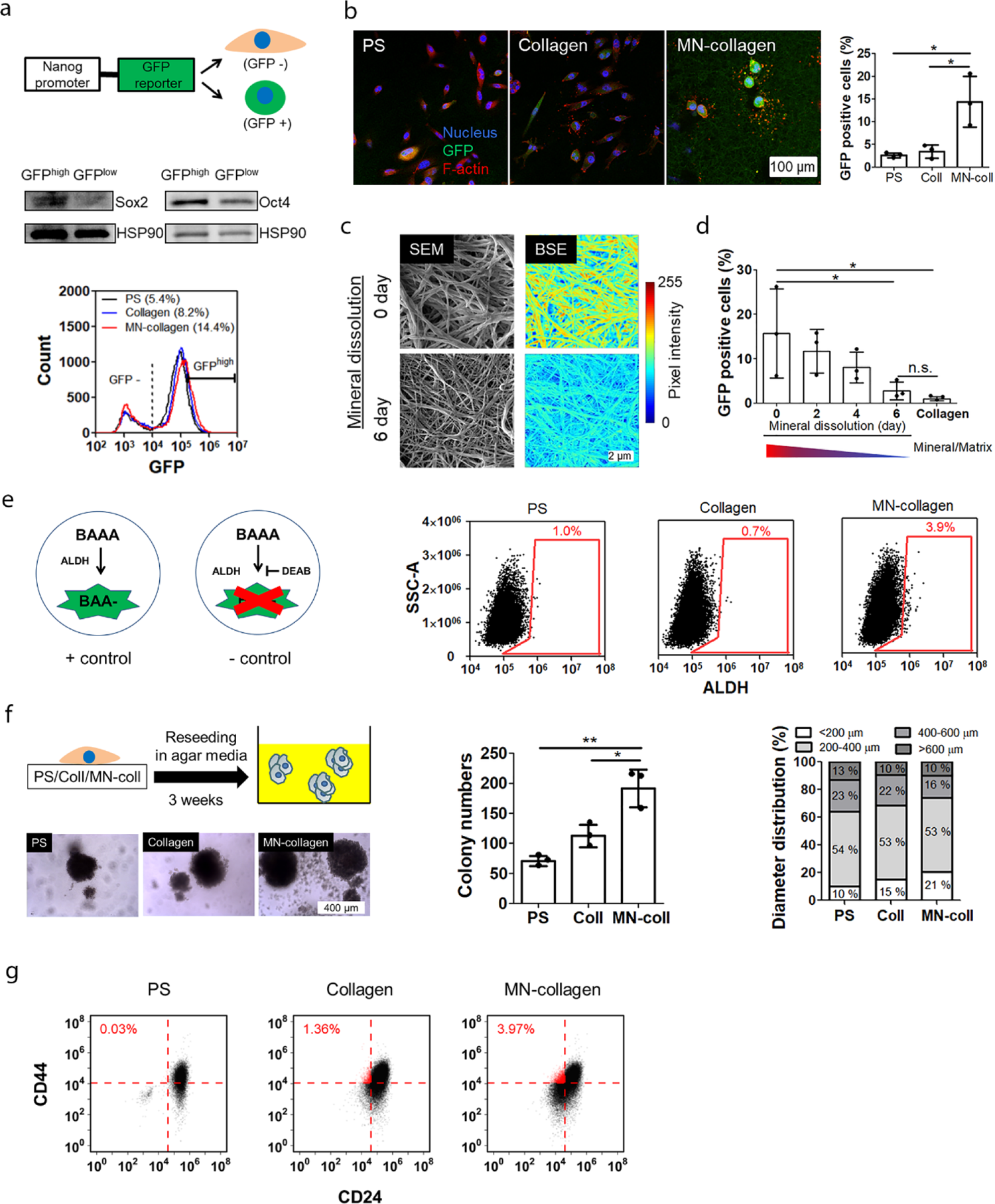
Collagen mineralization induces a stem-like phenotype in breast cancer cells. **a**, Schematic of the Nanog-MDA-MB231 reporter cell line in which increased GFP expression indicates a more stem-like phenotype (top). Western Blot analysis of Sox2 and Oct4 levels after FACS sorting of Nanog-MDA-MB231 reporter cells into GFP^high^ and GFP^low^ subpopulations (middle). Flow cytometry analysis of Nanog-MDA-MB231 reporter cells after 7 d of culture on polystyrene (PS), collagen, or mineralized collagen (bottom). **b,** Representative confocal micrographs and corresponding quantification of GFP positive cells after 7 d of culture on the different substrates (*n* = 3). Scale bar = 100 µm. **c**, **d**, Representative SEM and BSE (Pseudo-colored) images of mineralized collagen at different time point of dissolution (c) and confocal image analysis of GFP positive cells as a function of collagen mineral content (*n* = 3) (d). Mineral content of substrates was controlled by dissolution of hydroxyapatite over several days. One-way ANOVA post-hoc Dunnett’s multiple comparison test. **e**, Analysis of aldehyde dehydrogenase (ALDH) activity in parental MDA-MB231 cells. Aminoacetaldehyde (BAAA) is converted by ALDH into BODIPY-aminoacetate (BAA) that leads to increased intracellular fluorescence. Diethylaminobenzaldehyde (DEAB) inhibits ALDH and served as negative control. **f**, Soft agar colony formation after preculture of parental MDA-MB231 on the different substrates. Scale bar = 400 µm. Quantification of the number and size distribution of colonies larger than 100 µm in diameter (*n* = 3). **g,** Co-expression of CD44 and CD24 in MCF7 after 7 d of culture. The percentage of CD44^+^/CD24^-^ cells is indicated at the upper left of each panel. *: *P* < 0.05, **: *P* < 0.01.

To determine whether our findings were limited to MDA-MB231 cells, we also tested BoM-1833 but given the intrinsically high proliferative capacity of these cells their fraction of ALDH positive cells was negligible regardless of cell culture substrate (Supplementary Fig. 9). In contrast, MCF7 increased their stem-like properties following culture on mineralized collagen as demonstrated by a higher number of CD44^+^/CD24^-^ cells (Fig. 3g). For MCF7, CD44^+^/CD24^-^ represent appropriate stemness markers and characterize a population with more mesenchymal characteristics^62^ while ALDH is better suited for MDA-MB231 and BoM-1833 as almost all cells in these two lines are CD44^+^/CD24^-^^62^. Consistent with a more mesenchymal phenotype, MCF7 cells cultured on mineralized collagen also formed fewer cell-cell adhesions and had reduced E-cadherin levels (Supplementary Fig. 10).

### Mineralized collagen regulates breast cancer cell phenotype by altering mechanosignaling

Increased cell adhesion forces are drivers of tumor cell growth and typically increased on stiffer matrices^63^. While mineralization increases collagen stiffness, it simultaneously reduces collagen strain-stiffening and stress relaxation resulting in decreased rather than increased cell adhesion forces with consequential changes in cell spreading and collagen network remodeling (Fig. 1h, 2a, b)^19, 20, 64^. Because of these connections, we hypothesized that mineral-induced changes of mechanosignaling may be responsible for the above-described changes in tumor cell phenotype. To define which role mineralization-dependent changes in mechanotransduction play in regulating tumor cell growth, we cultured MDA-MB231 on collagen and mineralized collagen in the presence and absence of β1-integrin blocking antibodies and pharmacological inhibitors of FAK (FAK inhibitor 14), ROCK (Y27632), and PI3K (LY294002) (Fig. 4a). Treatment with these reagents reduced tumor cell growth on collagen as expected but had no effect on mineralized collagen suggesting that tumor cell mechanosignaling may be reduced on mineralized collagen. Tumor cells interacting with collagen have been described to increase their adhesion forces by aligning collagen fibers in a process that activates mechanosignaling via a positive feedback mechanism^41^. To test whether collagen mineralization interferes with this process by preventing cell-mediated collagen network remodeling and strain-stiffening as described above (Fig. 1h, 2b) and thus, reciprocal activation of cellular adhesion forces^20^, we performed traction force microscopy. Indeed, consistent with the limited effect of mechanosignaling inhibitors on tumor growth, MDA-MB231 pre-cultured on mineralized collagen exhibited reduced cellular traction forces relative to their counterparts precultured on collagen or PS (Fig. 4b, Supplementary Fig. 11) and sorted GFP^high^ cells exhibited similarly reduced traction forces relative to GFP^low^ cells (Fig. 4c). Consistent with these differences, MDA-MB231 cultured on mineralized versus control collagen adjusted their expression of genes related to cytoskeletal remodeling including *NEDD9* (a focal adhesion protein regulating cell attachment and the cell cycle), *KRT19* (intermediate filament proteins keratin), *S100A4* (encoding S100 proteins that regulate cell cycle progression and differentiation), and *FN1* (encoding fibronectin, which broadly regulates adhesion, but also dormancy^65^ (Supplementary Fig. 4a, c).

**Figure 4:**
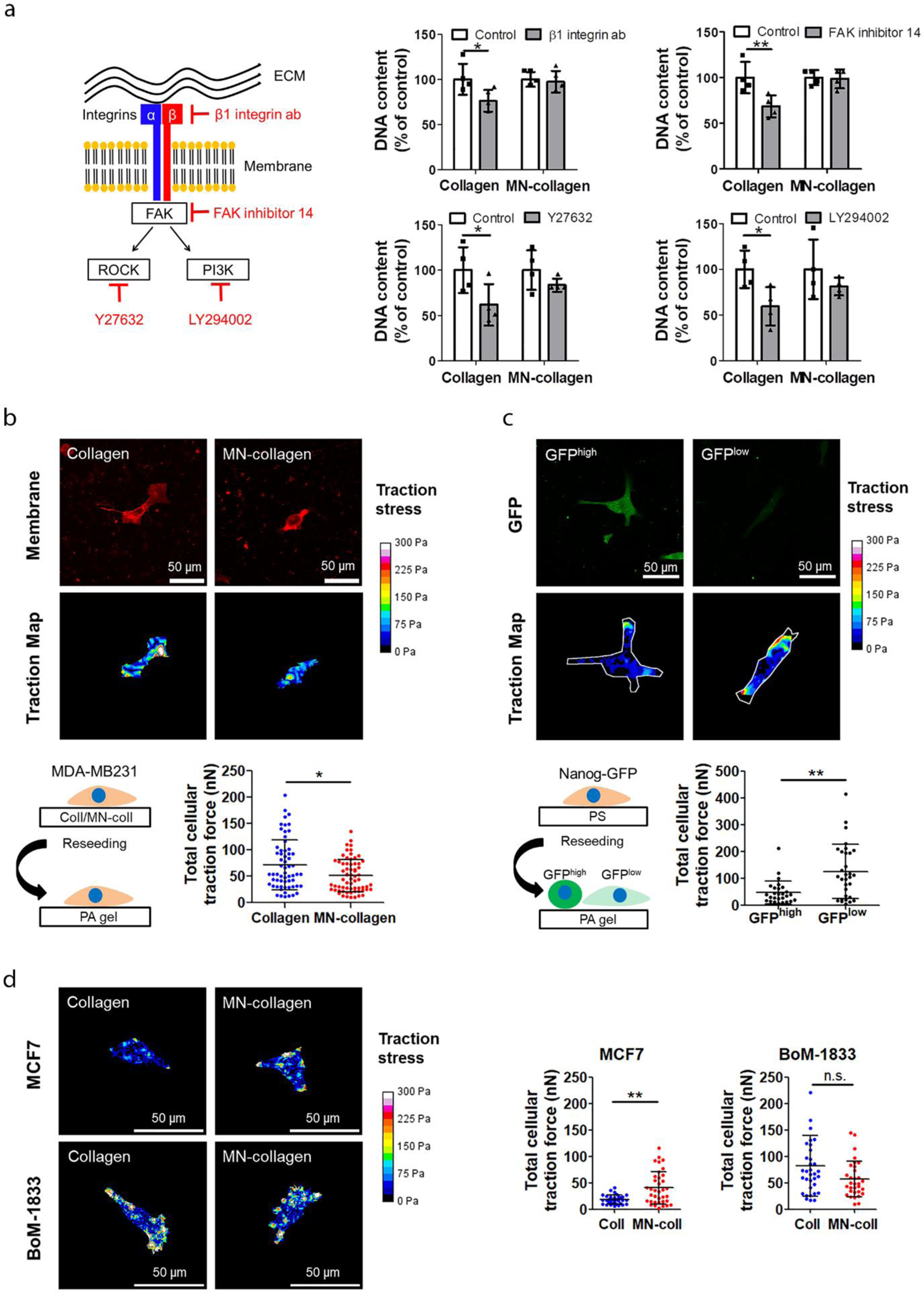
Collagen mineralization inhibits breast cancer cell mechanotransduction. **a,** Schematic of integrin-mediated mechanosignaling and effect of different mechanosignaling inhibitors on the growth of MDA-MB231 cells cultured on collagen and mineralized collagen (*n* = 4). **b,** Traction force microscopy of MDA-MB231 cells following preculture on collagen and mineralized collagen. Representative images of MemGlow 590-labeled cells (top), traction maps (middle), and corresponding quantification of traction forces (bottom). **c,** Traction force microscopy of GFP^high^ and GFP^low^ MDA-MB231 cells. Representative fluorescence images (top), traction maps (middle), and corresponding quantification of traction forces of Nanog-GFP cells. White lines on traction maps indicate contour of cell. **d** Traction force microscopy of MCF7 and BoM-1833 cells following preculture on collagen and mineralized collagen. Representative images of traction maps and corresponding quantification of traction forces (bottom). White lines on traction maps indicate contour of cell. Scale bar = 50 µm. *P* < 0.05 by Mann-Whitney U-test. *: *P* < 0.05, **: *P* < 0.01.

Interestingly, a similar experiment performed with MCF7 cells revealed that collagen mineralization increases rather than decreases MCF7 traction forces consistent with their increased mesenchymal properties (Fig. 4d, Supplementary Fig. 11). However, the absolute traction forces of MCF7 precultured on collagen were significantly lower than those of MDA-MB231. In contrast, traction forces of MCF7 precultured on mineralized collagen were similar to those of MDA-MB231 possibly suggesting that collagen mineralization leads to an adjustment of cell traction forces that are favorable for survival in bone. Indeed, the traction forces of BoM-1833, which were isolated from a bone metastasis, were similar to traction forces of MDA-MB231 and MCF7 cultured on mineralized collagen but did not differ when these cells were cultured on any of the other substrates (Fig. 4d, Supplementary Fig. 11). Collectively, these results suggest that collagen mineralization regulates the phenotype of breast cancer cells by inhibiting cell-mediated collagen remodeling that would otherwise increase growth by activating mechanosignaling.

### Mineral content in native bone matrix regulates breast cancer cell phenotype similar to mineralized collagen

In addition to collagen type I, the bone ECM contains other proteins (collectively referred to as noncollagenous proteins or NCPs) that may impact the response of tumor cells to mineral^66^. Additionally, the microfabricated collagen substrates described above are not suitable to validate *in vitro* results in a xenograft setting. To circumvent both limitations, we generated millimeter-sized physiological bone matrices in which we selectively controlled mineral content by adapting previously established protocols^67, 68^. We chose to prepare these bone matrices from trabecular bone given that it is a preferred site for breast cancer metastasis relative to long bones^69^. First, bone plugs were harvested from neonatal bovine femurs and fully decellularized to prepare scaffolds that contained all organic and inorganic matrix components including mineral but were devoid of cells (termed decellularized scaffolds [DC] from here on). Next, DC scaffolds were demineralized with ethylenediaminetetraacetic acid (EDTA) resulting in decellularized and demineralized (DCDM) scaffolds (Fig. 5a). Adequate decellularization was verified histologically and by DNA assay^70^ (Fig. 5b). Demineralization of DCDM was confirmed by nano-CT, BSE-SEM, and FT-IR, while SEM, second harmonic generation (SHG) microscopy, and histological analysis confirmed that scaffold architecture and collagen fibrillar structure were not affected by the demineralization procedure (Fig. 5c, Supplementary Fig. 12a, b, Supplementary Fig. 13). SEM analysis also indicated that the average diameter of collagen and mineralized collagen fibers in these scaffolds was comparable to the fiber diameter in our synthetic substrates, but more variable (Fig. 5c (ii)). Moreover, the pXRD patterns of DC scaffolds as well as their mineral to matrix ratio were comparable to physiological bone (and mineralized collagen) (Supplementary Fig. 1; mineralized collagen: approx. 4; human bone: 3-6)^26^ suggesting that the decellularization step had no effect on the mineral phase (Fig. 5d, e). Crystallinity of the HA in the DC scaffolds was also comparable to naïve bone but reduced relative to HA in mineralized collagen or pure HA, a difference that may be explained by the young age of the animals from which the bone was harvested (Fig. 5f).

**Figure 5:**
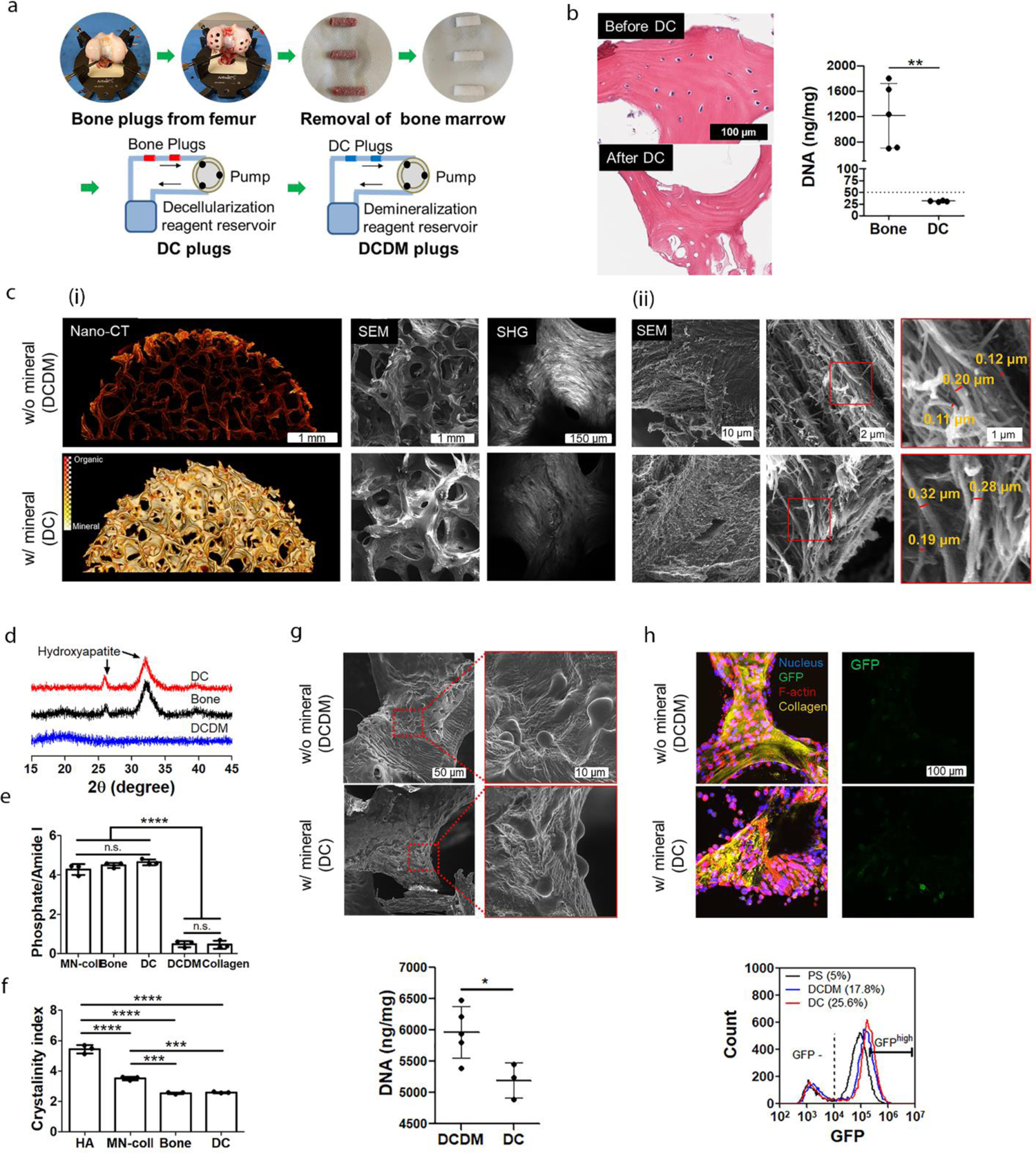
Physiological bone scaffolds to test the effect of bone mineral on the stem-like phenotype of MDA-MB231. **a,** Schematic visualizing the preparation of decellularized (DC) and decellularized/demineralized (DCDM) bone scaffolds from 1-2 cm long, 6mm diameter neonatal bovine femur bone plugs. **b**, H&E staining and DNA quantification of scaffolds after DC process (Bone: *n* = 5, DC: *n* = 3). Scale bar = 100 µm. **c**, Representative nano-CT, SEM and second harmonic generation (SHG) images (i) and SEM and fiber diameter analysis (ii) of scaffolds after DCDM treatment. Scale bar = 1 mm (Nano-CT and SEM) and 150 µm (SHG) (c(i)), and 10 µm (left), 2 µm (middle), and 1 µm (right) (SEM) (c(ii)). **d-f**, pXRD analysis of mineral phase **(d)** and FT-IR analysis of mineral to matrix ratio (*n* = 3) **(e)** and cyrstallinity index (*n* = 3**) (f)** after DC and DCDM process. Commercially available hydroxyapatite (HA) served as control. **g**, SEM images and DNA quantification of MDA-MB231 cells cultured on the different scaffolds (DCDM: *n* = 5, DC: *n* = 3). Scale bar = 50 µm (left) and 10 µm (right). **h**, Confocal images and flow cytometry analysis of Nanog-GFP cells cultured on the different scaffolds. Scale bar = 100 µm. *: *P* < 0.05, **: *P* < 0.01, ***: *P* < 0.001, ****: *P* < 0.0001.

After characterizing the bone scaffolds and confirming by SEM that MDA-MB231 cells adhered to and grew within the 3D scaffolds (Fig. 5g), we asked whether proliferation and stem-like behavior of breast cancer cells were altered by the different scaffold conditions. DNA analysis identified that cell growth within DC scaffolds was lower than that within DCDM scaffolds, supporting that the presence of mineral reduces breast cancer cell growth in these systems (Fig. 5g). Confocal imaging and flow cytometry of Nanog-GFP MDA-MB231 showed that breast cancer cells increased GFP expression and assumed more stem-like phenotypes when grown within mineral-containing DC scaffolds relative to PS and DCDM scaffolds (Fig. 5h). Taken together, these results indicate that the presence of mineral alters breast cancer cell behavior in physiological bone scaffolds similarly to the synthetic mineralized collagen scaffolds and thus, validate the broader significance of mineral to regulating tumor cell phenotypes.

### Bone mineral affects breast cancer progression *in vivo*

To test the relevance of bone matrix mineralization to tumor growth *in vivo*, decellularized physiological bone matrices with (DC) and without mineral (DCDM) (Fig. 5a) were seeded with luciferase-expressing MDA-MB231s, xenografted into female, athymic Nude-*Foxn1^nu^* mice, and then tracked longitudinally by bioluminescence (BLI) imaging (Fig. 6a). Tumor cells interacted with trabecular surfaces of both scaffold types but grew significantly more on matrices devoid of mineral (Fig. 6b, c). Accordingly, co-immunostaining against human vimentin and Ki67 confirmed that tumors in mineral-containing DC scaffolds contained fewer human tumor cells than their DCDM counterparts and that tumor cells were less proliferative relative to tumor cells growing in DC scaffolds (Fig. 6d). These results imply that mineralized bone matrix not only reduces tumor cell proliferation *in vitro*, but also *in vivo*.

**Figure 6:**
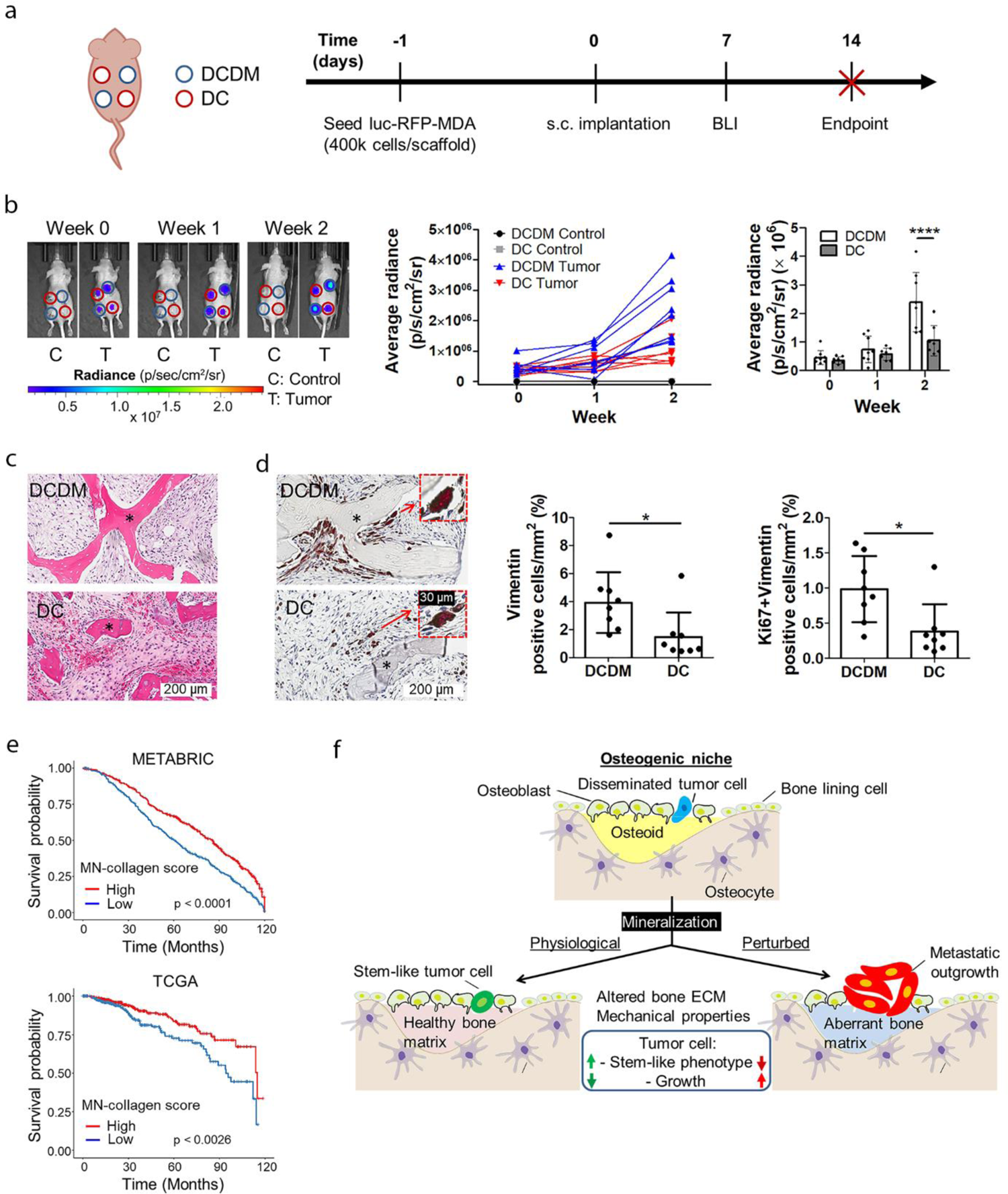
Mineralized bone matrix reduces tumor growth *in vivo,* and mineral-induced gene expression signature correlates with improved patient prognosis. **a,** Time course of xenograft experiment involving implantation of MDA-MB231-seeded DCDM and DC scaffolds into female nude mice. **b**, Representative bioluminescence intensity (BLI) images and BLI quantification of tumor growth on the different bone scaffolds (DCDM: *n* = 8, DC: *n* = 7). **c,d,** Representative H&E images (c) and IHC images and analysis of vimentin and vimentin/Ki67 positive cells (*n* = 8) (d) in implanted bone scaffolds. Scale bar = 200 µm and 30 µm (inset). * = scaffold. **e,** Overall survival of breast cancer patients scoring high or low for expression of mineral-induced gene signatures (Table 2) using the METABRIC and TCGA cohorts. **f,** Proposed relationship between bone ECM and breast cancer cell phenotype. Our results suggest that mineralization of collagen type I-rich osteoid in osteogenic niches induces stem-like, less-proliferative phenotypes in breast cancer cells. As bone matrix mineralization changes (e.g. due to aging etc.), tumor cell growth is activated possibly due to bone matrix-dependent changes in tumor cell mechanosignaling. *: *P* < 0.05, ****: *P* < 0.0001.

**Table 2.** Gene signature of MDA-MB231 cultured on mineralized collagen.

| Mineralized Collagen Gene Signature |  |  |  |
| --- | --- | --- | --- |
| ABCG1 | CHI3L2 | LINC01607 | SPINK4 |
| ABHD11-AS1 | CRIP2 | LIPG | STEAP4 |
| AC018816.1 | CSF2 | LRRK2 | TCP11L2 |
| AC024575.1 | CSRP3 | MEF2C | TFF1 |
| AC026403.1 | DEPP1 | MUC12 | TFF2 |
| AC095055.1 | DHRS9 | NANOS3 | TFF3 |
| ACOT1 | EGR3 | NEURL1 | TNFSF10 |
| ADM2 | EPHB6 | NFE2 | TPPP |
| AK8 | FAM20A | NNMT | TPPP3 |
| AL161787.1 | FBN2 | NR4A1 | TRBC1 |
| AL355312.3 | FN1 | NR4A2 | TRBV30 |
| AL356019.2 | FOS | NUPR1 | TRIB3 |
| AP000808.1 | FOSB | P2RX1 | TRPV6 |
| AQP1 | FOXS1 | PCDHGA7 | TSC22D3 |
| ATF3 | GAS7 | PHOSPHO1 | TSPAN8 |
| ATP6V0D2 | GKN2 | PRSS1 | TYMP |
| B4GALNT2 | HECW2-AS1 | PRSS2 | VLDLR-AS1 |
| B4GALNT3 | HPGD | PTGS2 | VWA1 |
| BMF | HSD3B1 | PTHLH | WFDC10B |
| C4B | IGFL2-AS1 | PTPRR | WFDC3 |
| CA14 | KLHL24 | RRAD | ZBTB20 |
| CD52 | KLRC2 | SERPINA5 | ZDHHC11B |
| CDC42EP5 | KPNA7 | SGK1 | ZNF474 |
| CDHR5 | KRT33B | SLC43A2 |  |
| CHAC1 | LINC00607 | SMIM1 |  |

To evaluate the clinical relevance of our findings, we examined whether the gene expression signature induced by culturing tumor cells on mineralized vs. control collagen was predictive of overall patient survival and bone metastasis-free survival. Indeed, upregulation of mineral-induced genes was associated with better patient prognosis in the METABRIC and TCGA cohorts consistent with our *in vitro* findings that matrix mineralization induces a more latent, less proliferative phenotype (Fig. 6e) and that tumor cells implanted on DCDM scaffolds grew more than their counterparts implanted on DC scaffolds. Analysis of cancer subtypes additionally showed that patients with luminal breast cancer, whose tumors primarily metastasize to bone, benefitted from upregulation of the mineral-induced gene signature (Table 2) whereas this was not the case for patients with basal breast cancer whose cancers preferentially metastasize to visceral organs (Supplementary Fig. 14)^71^. Using two published breast cancer cohorts with clinical data (Study ID: *GEO2603, GSE2034*) we further validated that patients whose tumors scored high for our mineralized collagen gene signature had improved bone-metastasis free survival relative to patients whose tumors scored low for this signature (Supplementary Fig. 15). While this analysis used primary tumor specimens rather than specimens collected from bone metastatic tumors, these results suggest that gene expression profiles induced by mineralized bone matrix correlate with increased patient survival possibly by selecting for less proliferative phenotypes. More broadly, these results indicate that bone matrix is a critical factor in regulating breast cancer cell fate.

## Discussion

Disseminating tumor cells target osteogenic niches within the skeleton^9, 10^ but how mineralization of collagen type-I, a hallmark of new bone formation, regulates the phenotype of breast cancer cells remains unexplored. Our results suggest that physiological collagen mineralization induces a less proliferative, more stem-like phenotype in breast cancer cells while perturbed mineralization activates tumor cells to proliferate and form larger tumors (Fig. 6f). These findings could help explain why breast cancer patients with decreased bone mineral density have a higher risk of developing bone metastasis^12, 27^.

Although ECM is known to regulate various aspects of breast cancer including metastasis,^72, 73^ its role in bone metastasis is less clear as studies in the skeleton are hindered by limited imaging modalities and difficulty of analyzing cells in this context. *In vitro* models can circumvent some of these limitations by allowing control over bone ECM materials properties. Here, we used biofunctional materials systems incorporating the basic building blocks of native bone ECM, hydroxyapatite-reinforced collagen fibrils ^18, 33^. With these models we were able to mimic defining materials properties of bone ECM across multiple length scales including bone’s hierarchical structure, ultrastructural arrangement, mineral-to-matrix ratio, and mineral crystallinity. Leveraging these models, we found that collagen mineralization suppresses cancer cell proliferation *in vitro* and tumor growth *in vivo*, and that mineral-induced changes in tumor stem-like properties may play a role in this process. These results suggest that osteogenic niches may induce quiescence in breast cancer cells due in part to collagen mineralization, giving new context to previous results showing that tumor cells assume a more stem-like phenotype following bone colonization^55^.

While matrix mechanical properties are widely accepted to affect tumorigenesis, the effect of bone matrix mineralization on tumor cell mechanosignaling is poorly understood due in part to a lack of model systems that mimic the viscoelastic properties of bone more appropriately than the linearly elastic substrates that are routinely used for mechanosignaling studies. Collagen type I, the primary organic bone matrix component, exhibits non-linear elasticity and viscoelasticity^74^. Both properties can affect cell behavior^21^ but are impacted by mineralization^19^. More specifically, tumor cells can activate and reinforce mechanosignaling by strain-stiffening collagen^41^, a process that is reduced by collagen mineralization as shown here. On the other hand, stress relaxation occurs more slowly in mineralized collagen than collagen^20^. Our data suggest that these changes have functional consequences as cancer cells adjust their traction forces following interactions with mineralized versus non-mineralized collagen. In MDA-MB231, these variations in traction forces manifested in a lack of response to pharmacological inhibitors of key nodes of mechanotransduction and correlated with reduced traction forces in more stem-like GFP^high^ versus GFP^low^ cells. Importantly, this correlation extended to an induction of phenotypic changes including upregulation of stem cell markers and functions, reduced cell proliferation, and increased anchorage-independent survival.

Our findings contradict the conventional assumption that bone stimulates malignancy by increasing mechanosignaling due to its increased rigidity and underline the importance of choosing the correct material systems for mechanistic studies of tumor-ECM interactions. Indeed, our results corroborate preclinical and clinical evidence indicating that reduced bone mineral density (e.g. due to Vit. D deficiency) correlates with increased risk and outgrowth of bone metastases^15, 16, 31^. Future studies will need to elucidate how mineralization-dependent changes of tumor cell phenotype affect other signaling mechanisms. For example, tumor cells can control entry into and exit from dormancy by depositing and responding to their own matrix including fibronectin^65, 75^, and altered cell-ECM interactions, in turn, can regulate cellular responses to soluble factors^65^. The fibronectin-binding integrins α_5_β_1_ and α_v_β_3_, for example, affect transforming growth factor-β (TGF-β) signaling via receptor cross-talk^76, 77^, a process that has been shown to modulate bone metastatic progression^13, 78^. Hence, future work should explore how interactions with mineralized collagen affect tumor cell deposition of fibronectin and, consequently, the expression of known dormancy markers such as NR2F1, which can regulate tumor cell stemness via Sox-2 and Nanog^79^. The influence of other nanoscale changes caused by mineralization, such as altered surface topography or porosity, should also be considered.

Although the focus of this study was on tumor cells, many other cell types residing in the skeleton can equally respond to bone matrix changes (e.g. endothelial cells, mesenchymal stromal cells, osteoblasts, osteocytes, immune cells)^9, 13, 80, 81^ and, vice versa, may alter bone matrix in response to tumor cell-secreted systemic or paracrine signals. Accordingly, bone matrix mineralization by osteoblasts correlated with decreased tumor cell growth in our experiments whereas inhibition of osteogenic differentiation and thus, matrix mineralization caused an opposite effect. While these data were supportive of our overall conclusion, we cannot exclude matrix-independent effects. For example, mineralizing, mature osteoblasts secrete factors (e.g. TGFβ2 or bone morphogenetic proteins 4 and 7 (BMP4/7)) that can induce tumor cell quiescence independent of matrix^82^, a possibility that should be tested with appropriate control experiments. Fluctuations in nutrient and oxygen supply or other metabolic constraints could further affect these responses. Future studies will need to increase the complexity of the described culture models to better understand how matrix mineralization impacts the early stages of bone metastasis in multivariate settings.

While we have validated the *in vivo* relevance of our findings with animal studies and with computational analysis of patient data, the model systems described herein have some limitations. For example, they did not consider bone-resident cells and the decellularized bone scaffolds were derived from neonatal bovine bone. The biomechanical properties of human bone and collagen are conserved between species, but it is likely that additional parameters influence bone matrix properties and thus, cancer cell response to mineralization. For example, age, gender, lifestyle, and prior treatment with chemotherapy are known regulators of bone material properties and health^23, 24, 28, 49, 83^. Future studies using bone specimens harvested from patients will help evaluate these connections and could be combined with matched bone-resident cells and organoid cultures to directly test patient-specific responses to bone matrix properties. Furthermore, bone mineralization is a dynamic process, and it is possible that tumor cell response is dependent on the time scales at which this process occurs. The mineralized substrates utilized here were prefabricated before seeding cells, but it is also possible to mineralize collagen *in situ*^81^. Finally, the flank implantation model we utilized allowed for precise control of bone matrix mineral content independent of other parameters, which is not readily possible in the skeleton itself. Biomaterials scaffolds can influence tumor growth and metastasis by altering the recruitment of immune and other stromal cells^84, 85^, and while we observed infiltration by resident cells, the mineral-induced less proliferative tumor cell phenotype was maintained. How the phenotype of recruited cells may differ in response to matrix mineral content and how such changes regulate tumor cell phenotype will need to be considered in future work.

In conclusion, our findings suggest that bone matrix changes are not only a collateral consequence of osteolytic degradation during late-stage metastasis but may also occur in osteogenic niches with potential effects on the early-stage development of bone metastases. The finding that bone matrix mineral content inhibits breast cancer growth motivates the clinical use of methods to increase or maintain physiological bone matrix mineralization, for example, by therapies that promote bone formation in settings of mechanical loading/exercise. Bisphosphonates are an alternative strategy to maintain bone mineral density and are routinely used as adjuvant therapy for breast cancer patients to reduce skeletal-related events and hypercalcemia. However, these drugs are only partially effective in preventing bone metastasis^86, 87^. As bisphosphonates primarily prevent bone resorption rather than encourage new bone formation and thus, matrix mineralization, our results could help explain the limited success of bisphosphonates in prevention settings. Collectively, our results motivate a more wholistic approach to modeling the bone ECM in future studies of bone metastasis and could yield insights that advance treatment options for breast cancer patients with advanced disease.

## Methods

### Fabrication of mineralized collagen

Mineralized collagen substrates for cell culture were fabricated using an adapted polymer-induced liquid-precursor (PILP) process (Fig. 1c) as previously described^20, 37^. Briefly, poly(dimethylsiloxane) (PDMS) (Dow Corning, US) microwells (diameter: 4 mm, height: 250 µm) were prepared on a larger, circular 8 mm PDMS base that enables face-down flotation of the devices necessary for intrafibrillar mineralization rather than sedimentation of mineral as further described below. Prior to casting collagen, the inner surface of the microwell was treated with oxygen plasma and 1% polyethyleneimine (PEI) (Sigma-Aldrich, US) and 0.1% glutaraldehyde (GA) (Thermo Fisher Scientific, US) to allow for subsequent covalent binding of pH-adjusted rat tail collagen, type I (Corning, US) (1.5 mg/mL). Mineralization was accomplished using a solution of 62.5 mg/mL of polyaspartic acid (PAA) (MW = 27k Da, Alamanda Polymers, US), 1.67 mM CaCl_2_ (Thermo Fisher Scientific, US) and 1 mM (NH_4_)_2_HPO_4_ (Sigma-Aldrich, US) in 0.85 × phosphate buffered saline (PBS). To prevent precipitation of mineral on top rather than within collagen fibrils, substrates were incubated in the mineralization solution upside down using a humidified chamber for 1 day at 37 °C. Collagen control substrates were fabricated similarly but immersed in PBS solution rather than mineralization solution. To adjust mineral content via controlled dissolution, fully mineralized collagen was incubated in 20 mM HEPES buffer (pH = 7.4) at ambient temperature for up to 6 days, changing HEPES buffer every 12 h.

### Characterization of mineralized collagen

For characterization, collagen substrates were prepared on PEI/GA treated 8 mm diameter glass coverslips adhered to a PDMS base. SEM (Mira3 LM, Tescan, Czech Republic) visualized fiber morphology and confirmed mineral formation using secondary electron (SEM) and backscattered electron (BSE) imaging modes. To this end, samples were dehydrated with a series of ethanol solutions and hexamethyldisiloxane and then carbon-coated (Desk II, Denton Vacuum, US). Changes in collagen fibril diameter were analyzed using the diameterJ function in ImageJ (NIH). Presence of mineral was detected by pseudocolor processing of BSE images using MATLAB (MathWorks, US). Mineral formation was also confirmed by Fourier transform infrared (FT-IR) spectrometry (Hyperion 2000/Tensor 27, Bruker, US). Samples were pelleted with potassium bromide (Thermo Fisher Scientific, US) and scanned in the range of 400 - 2000 cm^-1^. Mineral to matrix ratio was determined by calculating the ratio of phosphate peak area (907 - 1183 cm^-1^) and collagen amide I peak area (1580 - 1727 cm^-1^) after correcting the FT-IR spectra base line using spectroscopy software (OPUS, Bruker, US). The crystallinity index of mineral (CI; CI= (A_567_ + A_603_)/A_590_, where A*x* is the absorbance at wave number *x*) was determined using phosphate band splitting (See supplementary Fig. 1 and 3c). The phase of the formed mineral was determined using a powder X-ray diffractometer (XRD) (D8 advance ECO powder diffractometer, Bruker, US). Dried samples were mounted on a polymethyl methacrylate specimen holder and scanned in the range 2θ = 15 - 45° with a step size of 0.0195 degrees and Cu Ka radiation (λ = 1.54 Å). Mechanical properties of substrates were measured using a rheometer (DHR3, TA instruments, US) with 20 mm top- and bottom-plate geometry, a built-temperature and distance calibration. Collagen substrates were first cast and mineralized within a PDMS ring mounted onto 20 mm PEI-GA-coated coverslips. For rheological measurements, the PDMS ring was removed and the coverslips with the substrates were attached to the bottom-plate. Subsequently, a PEI-GA-coated coverslip was attached to the top-plate and the top-plate was lowered onto the sample at 500 µm distance between plates. In this configuration, the sample was incubated for 15 min at 37 °C in the presence of PBS before rheological measurements. For the storage (G’) and loss (G”) moduli, time sweeps were performed at 1% strain with an oscillation frequency of 2π rad/s. Strain stiffening was measured using strain sweep experiments with an oscillation frequency of 1 rad/s. Stress relaxation was measured using time sweeps experiments after applying 40 % strain with a rise time of 0.01 s.

### Cell culture and characterization of adhesion and proliferation

MDA-MB231 breast cancer cells (ATCC), MDA-MB231 expressing a Nanog-GFP reporter^56^ (Nanog-GFP), bone metastatic BoM1-2287 and BoM-1833 (kindly provided by Joan Massague), and MCF7 (ATCC) were routinely cultured in Minimum Essential Medium α (α-MEM) (Thermo Fisher Scientific, US) supplemented with 10% (v/v) fetal bovine serum (FBS) (Atlanta Biologicals, US) and 1% penicillin/streptomycin (P/S) (Thermo Fisher Scientific, US). Substrate-dependent differences of cell adhesion and growth were determined by fluorescence analysis of DNA content with the QuantiFluor dsDNA kit (Promega, US) after 5 h and 7 d of culture, respectively. For analysis of fibronectin effect on cell growth, substrates were incubated with PBS solution containing various concentrations of human plasma fibronectin (Thermo Fisher Scientific, US) for 2 h. Prior to cell seeding, fibronectin-adhered substrates were transferred to cell culture media containing 10% FBS and 1% P/S. For analysis of cell morphology after 5 h of adhesion, cells were fixed with 10% formaldehyde and stained with DAPI (Invitrogen, US) and Alexa Fluor 568 phalloidin (Thermo Fisher Scientific, US). Confocal images (710 LSM, Zeiss, Germany) were captured with a 40× water immersion objective at 2 µm step size. Cell morphology was analyzed using randomly selected images per sample (*n* = 4). At least 200 cells per condition were analyzed with built-in functions of Image J software for projected cell area and aspect ratio. Collagen structure was imaged in reflectance mode using a 488 nm laser. Cell proliferation was assessed after 7 days using Ki67 staining. Fixed samples were blocked with 1% BSA in PBS, stained using rabbit anti-human Ki67 monoclonal antibody (Cell Signaling Technologies, US), detected with goat anti-rabbit Alexa Fluor 568 secondary antibody (Thermo Fischber Scientific, US), and counterstained with DAPI. Ki67 levels per cell were analyzed from at least four randomly selected areas per sample (*n =* 4) and quantified using ImageJ. Cell proliferation was also measured after 4 days using a BrdU staining kit (Life Biotechnologies, US) according to manufacturer’s instructions. Images from six randomly selected areas per sample (*n* = 3) were captured with an epifluorescence microscope and BrdU positive cells were quantified using ImageJ.

### Experiments using human, bone marrow-derived mesenchymal stem cells

Human Bone-Marrow Derived MSCs (RoosterBio, US) were routinely cultured in RoosterNourish™-MSC media. MSCs were seeded on collagen-coated 18mm glass coverslips. Tumor conditioned media (TCM) was generated by culturing MDA-MB231 breast cancer cells (ATCC) until 90% confluency in DMEM/10%FBS/1%P/S, then media was replaced with serum-free media (DMEM, 1% P/S) for 24 h. Conditioned media was collected and concentrated 10-fold in an Amicon centrifugal filter unit (MWCO 3k Da, EMD Millipore) and subsequently diluted 5-fold with MSC media. MSCs were treated with TCM, osteogenic induction media (50 μM ascorbic acid, 0.1 μM dexamethasone, 10 mM β-glycerophosphate) and/or control DMEM media every 2-3 days. After 21 days, MSC mineral deposition was assessed via Alizarin Red S staining. Samples were fixed with 4% PFA for 15 min and incubated with 40 mM Alizarin Red S (VWR, US) for 30 min, followed by washing with distilled water. To assess MSC fibronectin deposition, fixed samples were blocked with 1% BSA in PBS, stained using mouse anti-human fibronectin monoclonal antibody (Sigma-Aldrich, US) and detected with goat anti-mouse Alexa Fluor 488 secondary antibody (Thermo Fisher Scientific, US). Cells were counterstained with DAPI (Invitrogen, US).

To assess changes in cancer cell proliferation after MSC pretreatment, MDA-MB231 cells were labeled with CellTracker™ Orange CMRA dye (Invitrogen, US) according to manufacturer’s instructions and seeded onto MSC cultures that were treated for 21 days. After 2 days of culture, samples were fixed with 4% PFA and counterstained with DAPI (Invitrogen, US). Confocal images (710 LSM, Zeiss, Germany) were captured with a 10× air objective from five randomly selected areas per sample (*n* = 3) and tumor cell number (per field of view) was quantified using ImageJ.

### Analysis of stem-like tumor cell phenotype

To validate the Nanog-GFP reporter cell line, cells were sorted into GFP^high^ and GFP^low^ populations using FACS. Sorted cells were lysed with RIPA buffer containing protease and phosphatase inhibitors (Thermo Fisher Scientific, US) and 1 mM phenylmethylsulfonyl fluoride (Calbiochem, US). All samples with equal amounts of protein were loaded on gels and separated by reducing SDS-PAGE and transferred to PVDF membranes (Bio-Rad, US). Membranes were blocked with 5% milk powder and incubated with rabbit anti-human Sox2 (Sigma-Aldrich, US), rabbit anti-human Oct4 (Millipore, US), and rabbit anti-human HSP90 (Santa Cruz, US) overnight. Primary antibodies were detected by a horseradish peroxidase (HRP)-conjugated anti-rabbit antibody using an ECL kit (Thermo Fisher Scientific, US) and imaged using ChemiDoc TM Touch Imaging System (Bio-Rad, US). Captured images were analyzed with Image Lab software (Bio-Rad, US). For analysis of matrix effect on GFP expression, Nanog-GFP cells were cultured on the different matrices and GFP levels per cell were analyzed from confocal images using the threshold tool in ImageJ. For flow cytometry Nanog-GFP cells cultured on tissue culture polystyrene (PS) were trypsinized, while cells cultured on collagen and mineralized collagen were isolated using collagenase, Type I (1 mg/mL in PBS, Worthington Biochemical, US). Suspended cells were passed through a 40 µm cell strainer, centrifuged, and resuspended in flow cytometry buffer containing propidium iodide (PI) (Invitrogen, US) for subsequent flow cytometry (BD Accuri C6 Plus, BD biosciences, US). Aldehyde dehydrogenase (ALDH) activity of cells was measured using an ALDEFLUOR kit (STEMCELLTechnologies, CA) according to manufacturer’s instructions. Cells were gated by FSC and SSC, then FSC-A and FSC-H signals were used for doublet exclusion. Live cells selected based on PI signals were used to identify cell populations expressing GFP from Nanog-GFP and ALDH from MDA-MB231. To analyze colony formation in soft agar, cells isolated from the different matrices (5 × 10^4^ cells) were suspended in a solution of 0.3% noble agar in Dulbecco’s Modified Eagle Medium (DMEM) with 10% FBS and 1% P/S and then plated on a layer of 0.5% noble agar in 6-well plates. After gelation, the plate was cultured in DMEM with 10% FBS and 1% P/S. After 21 days, cultures were incubated with media containing nitroblue tetrazolium (Biotium, US) overnight. Sphere formation was visualized using a ChemiDoc Touch Imaging System (Bio-Rad, US) and analyzed using the ImageJ particle analysis tool. Colonies over 100 µm of diameter were counted and measured for colony diameter distribution. For analysis of stemness and E-cadherin expression of MCF7 cells, MCF7 cells were cultured on the different matrices for 7 days. MCF7 cells were assessed for the CD44^+^/CD24^-^ stem-like fraction via flow cytometry using a BD Accuri C6 Plus Analyzer. Following trypsinization, cells were resuspended in FACS buffer (PBS containing 2.5% FBS, 2 mM ethylenediaminetetraacetic acid (EDTA)) at 10 × 10^6^ cells/mL followed by incubation with antibodies against human CD44 (APC-conjugated, Clone G44-26, 1:5, BD Biosciences, US) and CD24 (PE-Cy7-conjugated, Clone ML5, 1:20, BD Biosciences, US). Gates were determined using the isotype controls mouse anti-IgG2b κ (APC-conjugated, Clone 27-35, 1:5, BD Biosciences, US) and mouse anti-IgG2a κ (PE-Cy7-conjugated, Clone G155-178, 1:20, BD Biosciences, US). E-cadherin expression was analyzed from confocal images. Cells were stained using mouse anti-human E-cadherin (BD Biosciences, US) and detected with goat anti-mouse Alexa Fluor 568 secondary antibody (Thermo Fisher Scientific, US). E-cadherin levels per cell were analyzed from four randomly selected areas per sample (*n* = 3) and quantified using ImageJ.

### Analysis of mechanosignaling-mediated changes of cell growth

For analysis of mineralization-dependent changes of mechanosignaling, MDA-MB231 cells were cultured on PS, collagen and mineralized collagen in the presence and absence of a function-blocking β1 integrin antibody (2 µg/mL, Millipore, US) or pharmacological inhibitors of focal adhesion kinase (FAK) (FAK inhibitor 14, 5 µM, Tocris, UK), Rho-associated protein kinase (ROCK) (Y-27632, 25 µM, Tocris, UK), or phosphoinositide 3-kinase (PI3K) (LY 294002, 10 µM, Tocris, UK) for 4 days. Subsequently, cell growth was analyzed by measuring DNA content as described above.

### Analysis of cell traction forces

Polyacrylamide gels were prepared with a Young’s modulus of either 2.7 kPa for Nanog-GFP and 5 kPa for MDA-MB231 cells as previously described^88^. Briefly, 35 mm glass-bottom dishes (VWR, US) were cleaned with 0.1 N NaOH, followed by sequential surface treatment with 3-aminopropyl-trimethylsilane and 0.5% (v/v) glutaraldehyde in PBS. Dishes were washed with distilled water (dH_2_O) and allowed to air dry. Prepolymer solutions for gels were prepared by combining 40% acrylamide, 2% bisacrylamide, PBS, and 0.2 µm fluorescent beads (Thermo Fisher Scientific, US). 2.7 kPa and 5 kPa gels were prepared using 7.5%/0.035% and 7.5%/0.06% of acrylamide/bisacrylamide, respectively. Polymerization was initiated through the addition of ammonium persulfate and N,N,N,N-tetramethyl ethylenediamine. Droplets of the polymer solution were placed between activated dishes and a Sigmacote®-treated 8 mm coverslip (Sigma-Aldrich, US), and allowed to polymerize. Polyacrylamide gels were functionalized with rat tail type-I collagen (0.03 mg/mL) using an N6 succinimide ester crosslinker solution as previously described^89^. Gels were washed with PBS and cell media prior to cell seeding at 1 × 10^3^ cells/cm^2^.

Confocal microscopy (810 and 880 LSM, Zeiss, Germany) was used to image Nanog-GFP or MemGlow 590 (20 nM, Cytoskeleton, US) labeled MDA-MB231 cells using either 63× or 40×, 1.2NA C-Apochromat objectives. Cells were lysed by adding 2% sodium dodecyl-sulfate (SDS) to the dishes (final concentration of 1% SDS) and gels allowed to relax for 2 min. Images of the relaxed gels were then acquired at the same positions. Images of beads were sharpened, and stage drift corrected using ImageJ, and bead displacements and corresponding forces calculated using previously developed software and custom MatLAB® scripts^90^. Cellular traction forces were calculated using the Fourier transform traction cytometry (FTTC) method with a regularization parameter (λ) of 1 × 10^-9^. For MDA-MB231, MCF7, and the bone tropic MDA-MB231 subclone BoM-1833 traction forces of at least 10 cells were analyzed per gel for a minimum of 2 gels per condition. For Nanog-GFP, traction forces of at least 25 cells per condition were analyzed from 5 gels. Pairwise comparisons of total cellular traction forces were conducted using either a Mann-Whitney U-test or a Kruskal-Wallis test with Dunn’s multiple comparison correction when more than two conditions were being compared. For cellular traction forces when cells were precultured on different substrates, outliers were identified using the ROUT method with a Q value of 1%.

### RNA-sequencing (RNA-seq) and data analysis

Cells were collected using collagenase after 7 days of culture, and RNA was isolated from each condition using an RNA isolation kit (Qiagen, US) according to the manufacturer’s protocols (*n* = 4). Illumina TruSeq RNA stranded kit (Illumina, US) was used for library preparation, and samples were sequenced as 1 × 86 bp single-end reads on the NextSeq500 (Illumina, US), yielding on average 42 M reads per sample. Reads were then trimmed using Trim Galore (version 0.4.4; https://www.bioinformatics.babraham.ac.uk/projects/trim_galore/), and aligned to the human reference genome GRCh38 (ENSEMBL) using STAR (version 2.6.0a)^91^. Reads of genomic features were counted using featureCounts^92^, and differential gene expression determined using DESeq2^93^. Differentially expressed genes were defined as those with a log_2_-fold change greater than |1| and an adjusted p-value less than 0.05. For gene set enrichment analysis (GSEA), a ranked list was generated by taking the sign of the fold change multiplied by the -log_10_ of the adjusted p-value. The list was inputted to the GSEA Java applet (http://software.broadinstitute.org/gsea/index.jsp) using gene ontology biological processes gene sets from MSigDB v.6.2. Gene sets were considered significantly enriched with a p-value and FDR value < 0.05.

The relevance of gene expression changes induced by mineralized collagen to human breast cancer samples was computed using publicly available TCGA and METABRIC datasets. The 98-gene mineralized collagen gene signature was defined as transcripts with a log_2_-Fold Change ≥ 1 and a p-adjusted value ≤ 0.05. The transcriptional score based on supervised gene signature was computed across all samples using the Single Sample Gene Set Enrichment Analysis method (SSGSEA) from the *gsva* package. SSGSEA scores were utilized for clinical correlations with survival. In TCGA and METABRIC cohorts, sorted signature scores in the upper and lower tertiles were classified as high and low transcriptional signature, respectively. Kaplan-Meier survival analysis was then performed using coxph regression using the *survival* package in R to associate transcriptional signature score with overall survival as primary outcome.

To assess association of our mineralized collagen gene signature with bone-metastasis free survival two published breast cancer cohorts with clinical data (Study ID: *GEO2603, GSE2034*) were utilized. ComBat function from the sva library was used to batch correct the cohorts prior to the analysis. As the utilized published data were generated by microarray, which analyzes predefined transcripts/genes rather than the whole transcriptome, we were only able to map a subset of our RNA-Seq derived 98 gene signature to the microarray probe (GLP96) set in this cohort. Despite this technological limitation with the dataset, we were able to use 55 out of the 98 genes that mapped to the available probe set. This subset of 55 genes was used as the mineralized collagen gene signature to compute the transcriptional signature score using SSGSEA as above. Across the cohorts, sorted signature scores in the upper and lower quartiles were classified as high and low transcriptional signature, respectively. Kaplan-Meier survival analysis was then performed using coxph regression using the *survival* package in R to associate transcriptional signature score with bone metastasis free survival.

### Preparation and characterization of bone scaffolds

Trabecular bone biopsies were extracted from 1-3 day-old neonatal bovine distal femurs as previously described^67^. After rinsing with a high velocity stream of deionized water to remove marrow and debris, biopsies were sectioned into cylindrical scaffolds (6 mm-diameter and 1 mm-thickness). Scaffolds were incubated in an extraction buffer of 20 mM NaOH and 0.5% Triton X-100 in PBS at 37 °C to remove cells and cellular debris, then incubated in 20 U/mL DNase I to remove any latent DNA fragments (decellularized [DC] scaffolds). Bone mineral was removed from DC scaffolds with an overnight incubation in 9.5% EDTA (decellularized and demineralized [DCDM] scaffolds). All scaffolds were then washed 5 times with PBS to clear any residual material. Removal of cellular components was confirmed by H&E staining as well as DNA quantification. To confirm demineralization, scaffolds were scanned on a high resolution 3D X-ray microscope (Xradia Zeiss Versa XRM-520, Zeiss, Germany) at 60 kV/5W, with exposures of 1.5 s each, at a resolution of 3.53 microns/pixel, and false-colored based on the attenuation coefficient (Avizo, Thermo Fisher Scientific, US). To assess collagen content, Masson’s trichrome staining was performed on formalin fixed, paraffin-embedded sections according to standard protocols. Trabecular structure and collagen fibrillar structure in DC and DCDM scaffolds were determined by SEM while presence of mineral was determined by BSE. Collagen microstructure of DC and DCDM scaffolds was also measured by second harmonic generation (SHG) microscopy. XRD and FT-IR spectra were used to characterize mineral properties of scaffolds as described above. For detection of cell growth, MDA-MB231 cells were imaged by SEM and quantified by fluorescence analysis of DNA content with the QuantiFluor dsDNA kit (Promega, US). For detection of stem-like properties of cells in scaffolds, GFP expression of Nanog-GFP was imaged by confocal microscopy and quantified using flow cytometry.

### Animal Studies

Mouse studies were performed in accordance with Cornell University animal care guidelines and was approved by Cornell University’s Institutional Animal Care and Use Committee (IACUC). For detection of disseminated tumor cells relative to mineralizing bone surfaces, MDA-MB231 cells were labeled with fluorescent silica nanoparticles containing positively charged Cy5 dye (Cy5(+)-SNPs) as previously described^35^. Briefly, cells were labeled with 1 µM Cy5(+)-SNPs and resuspended in ice-cold PBS for mouse injections. Female athymic nude-Foxn1nu (Envigo, US) mice were injected intraperitoneally with calcein (15 mg/kg, pH 7.4 in PBS) at 5-6 weeks of age. One day later, labeled cells (1 × 10^5^ cells in 100 µL PBS) were injected into the left cardiac ventricle using ultrasound guidance (Vevo VisualSonics 2100, FUJIFILM VisualSonics, Canada). Injection into the systemic circulation was verified by bioluminescence imaging (IVIS Spectrum, Perkin Elmer, US). Tibiae were harvested after 2 days. Immediately following harvest, tibiae were fixed in ice-cold 4% paraformaldehyde (pH 7.4 in PBS) for 16 h. To detect tumor cells labeled with Cy5(+)-SNPs, tibiae were optically cleared using ethyl cinnamate following dehydration in a graded ethanol series^94^. Samples were imaged in ethyl cinnamate using a light sheet microscope (LaVision BioTec, Germany), using the 488 nm laser to detect calcein and the 640 nm laser to detect Cy5(+)-SNPs. Arivis Vision4D (Arivis, Germany) and ImageJ were used to process light sheet microscopy images.

For *in vivo* tumor formation, luciferase-expressing MDA-MB231 were seeded on bovine bone scaffolds (6 mm-diameter and 1 mm-thickness) in α-MEM with 10% FBS and 1% P/S (4 × 10^5^ cells/scaffold) and cultured at 37 °C in a 5% CO_2_ incubator for 24 h prior to implantation. Cell-laden scaffolds (decellularized [DC] vs decellularized and demineralized [DCDM]) were implanted subcutaneously into six- to eight-week-old female BALB/c athymic nude mice (Taconic Biosciences, US). Two scaffolds per condition were implanted into each mouse (*i.e.,* 4 scaffolds per mouse; *n*=4 mice; 8 scaffolds per condition). Bioluminescence images were taken once weekly 5 min following injection of D-luciferin (Gold Biotechnology, US) with an in vivo imaging system (IVIS Spectrum, Perkin Elmer, US). Outliers were determined using a Grubb’s test with a significance level of alpha = 0.0001. Explanted scaffolds were fixed with ice-cold 4% paraformaldehyde and decalcified with 10% EDTA. Paraffin sections were used for H&E staining as well as sequential double-immuno-staining with rabbit anti-human Ki67 (Cell Signaling Technologies, US) and rabbit anti-human vimentin (Thermo Fisher Scientific, US). First, Ki67 antibody was detected with an alkaline phosphatase (AP)-conjugated horse anti-rabbit antibody to stain red following incubation with AP substrate (Vector Laboratories, US). After blocking Ki67-stained samples with horse serum, human vimentin antibody was detected with a HRP-conjugated horse anti-rabbit antibody and diaminobenzidine (DAB) to produce a brown color (Thermo Fisher Scientific, US). All sections were counterstained with Mayer’s hematoxylin (Thermo Fisher Scientific, US) and imaged using a ScanScope slide scanner (Aperio CS2, Leica Biosystems, Germany) with a 40× objective. To quantify Ki67 and vimentin immunoreactivity, images of all IHC-stained sections were uploaded to QuPath v0.2.0^95^ and manually annotated to include cells and exclude trabeculae, lipid deposits, and extracellular debris. A QuPath script was written to quantify the total number of cells, total section area, and number of cells positive for vimentin and Ki67 per section area. Each data point represents the percentage of positive cells per mm^2^ as identified for the cross-sectional area of 1 implant; 8 implants were analyzed per condition.

### Statistical analysis

Unless otherwise indicated results are presented as the mean and standard deviation (SD) of at least 3 independent replicates per condition using Prism 8 software (GraphPad, US). Statistical differences were determined by Student’s unpaired *t*-test for two group comparisons, one-way analysis of variance (ANOVA) followed by Tukey’s post test for multiple group comparisons and two-way ANOVA with Sidak’s multiple comparison for multiple factors, unless otherwise mentioned. Two-tailed test with *P* < 0.05 were considered significant in all cases. Statistical considerations for analysis of cell traction forces and tumor growth *in vivo* are specified in the respective sections above.

## Acknowledgements

We thank all members of the Fischbach lab for valuable discussions of this research, the Wiesner group for providing fluorescent silica nanoparticles, Joe Kuo for help with illustrations, Lauren O’Keeffe for help with preparation of bone scaffolds, the Cornell Animal Health Diagnostic Core for paraffin embedding and sectioning, and the Cornell Center for Animal Resources and Education (CARE) staff for animal care. Financial support was provided by the Human Frontier Science Program (RGP0016/2017), the National Cancer Institute through the Center on the Physics of Cancer Metabolism (1U54CA210184), and NSF GRFP to M.A.W and A.A.S. (DGE-1650441). This work utilized the Cornell Center for Materials Research (CCMR), which is supported through the NSF MRSEC program (DMR-1719875), the Cornell NanoScale Science & Technology Facility (CNF), a member of the NSF-supported National Nanotechnology Coordinated Infrastructure (NNCI-2025233), and the Cornell University Biotechnology Resource Center (BRC) facilities, including Zeiss LSM 710 Confocal Microscope (NIH 1S10RR025502), Zeiss LSM880 confocal/multiphoton microscope (NYSTEM (C029155) and NIH (S10OD018516)), light sheet microscope (NIH S10OD023466), ZEISS/Xradia Versa 520 X-ray Microscope (NIH S10OD012287), IVIS Spectrum (NIH S10OD025049), and Genomics Facility (RRID:SCR_021727).

## Author contributions

S.C., M.A.W., L.A.E. and C.F. designed the project. S.C. and M.A.W. conducted most of the experiments. A.A.S. performed and analyzed FACS and RNA-seq. N.D.S performed and analyzed hMSC experiments. A.C. performed the animal study for light sheet microscopy. J.D. performed FACS. A.V. and O.E. analyzed RNA-seq data with SSGSEA method. S.L analyzed IHC images. Z.C, A.A.S., and M.P. conducted and analyzed TFM. S.C., M.A.W., L.A.E. and C.F. analyzed the data and wrote manuscript. All authors discussed the results and commented on the manuscript.

## Data availability

The main data supporting the results in this study are available within the paper and its Supplementary Information. All RNA sequence data generated in this study are available from the corresponding author upon reasonable request. Source data for the figures are provided with this paper.

## Code availability

QuPath script is available from the corresponding author upon request.

