## Supplemental figures for "Biofunctional matrix models reveal mineral-dependent mechanoregulation of bone metastatic breast cancer"

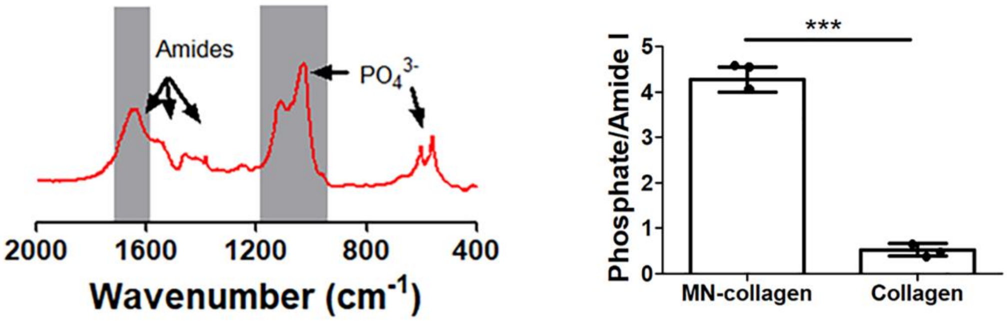


**Supplementary Figure 1:** Mineral to matrix ratio of mineralized collagen as calculated by dividing the phosphate peak area (907 - 1183 $cm^{-1}$) of FT-IR spectra by the amide I peak area (1580 - 1727 $cm^{-1}$) ($n$ = 3). ***: $P < 0.001$ by Student's unpaired $t$-test.

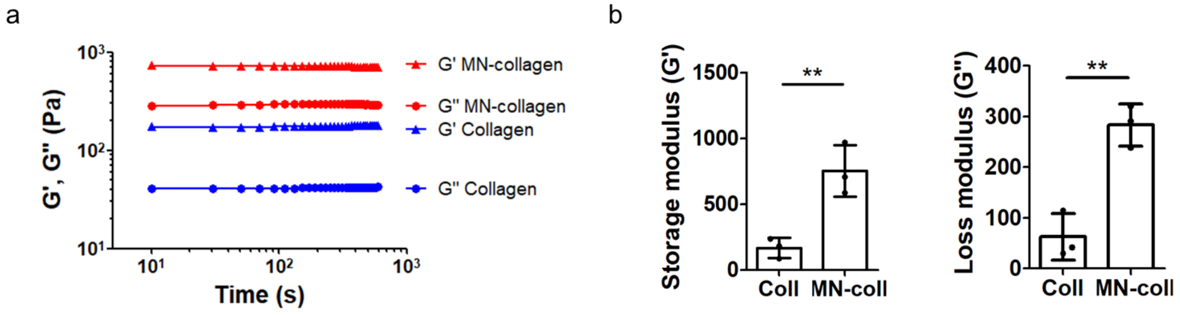


**Supplementary Figure 2: a, b,** Representative time sweeps (a) and quantification of storage (G′) and loss (G″) moduli ($n$ = 3) (b). **: $P < 0.01$ by Student's unpaired $t$-test.

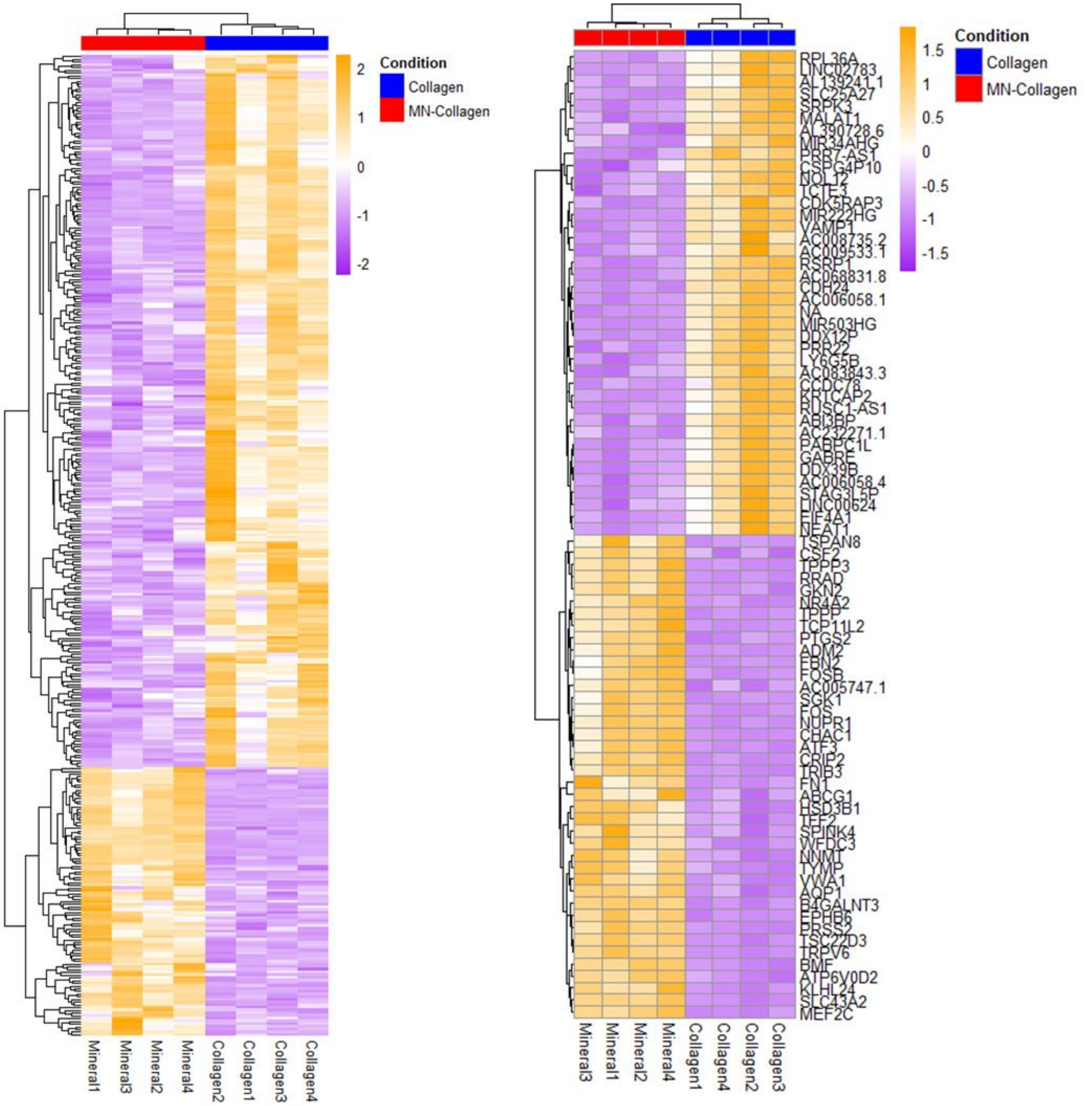


**Supplementary Figure 3:** Heatmap of differentially expressed genes (DEGs). DEGs were defined as the genes with FDR-adjusted p-values < 0.05 and $\log_2 FC > 1$. Relative expression levels of all DEGs (left) or top 50 genes (right) is split into collagen and mineralized collagen conditions.

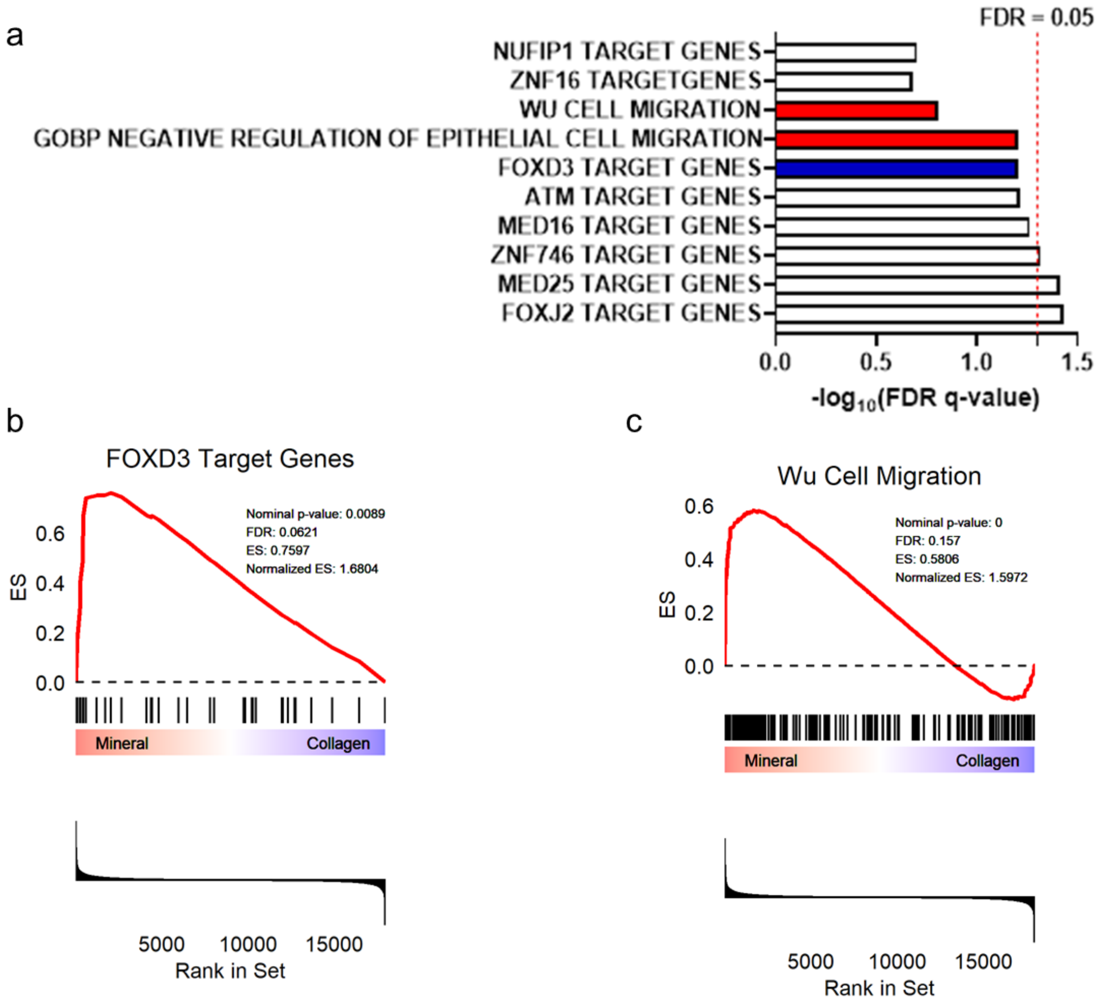


**Supplementary Figure 4: a**, Top 10 enrichment plot from gene set enrichment analysis (GSEA) indicates that biological processes associated with transcription and cell migration were overrepresented by collagen mineralization. Red labeled gene sets are associated with actin remodeling and migration curated from the Molecular Signatures Data Base (MSigDB), while blue labeled gene sets have been directly linked to stemness. The top 3 significant transcription factors (*FOXJ2*, *MED25*, *ZNF746*) do not have clearly defined functions beyond complexing with RNA polymerase II but *FOXD3* was also enriched albeit to a less significant extent (FDR q-value = 0.062). Dashed red line designates FDR q-value cutoff of 0.05. **b,** Enrichment plot for FOXD3 target genes as determined by GSEA. Normalized Enrichment Score (NES) = 1.68, nominal p-value = 0.009, FDR q-value = 0.062. **c,** Enrichment plot for the Wu Cell Migration gene signature from the MSigDB as determined by GSEA. Normalized Enrichment Score (NES) = 1.60, nominal p-value < 0.001, FDR q-value = 0.157. While cell migration was not a focus of our study, this gene set includes several genes directly related to cytoskeletal remodeling (e.g. *NEDD9*, *KRT19*, *S100A4*, and *FN1*).

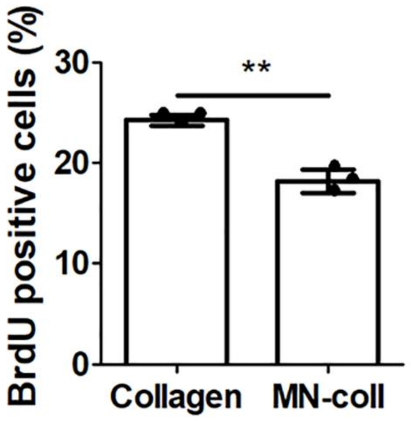


**Supplementary Figure 5:** BrdU image analysis and of MDA-MB231 cells cultured for 4 d on the different substrates ($n$ = 4). **: $P < 0.01$ by Student's unpaired *t*-test.

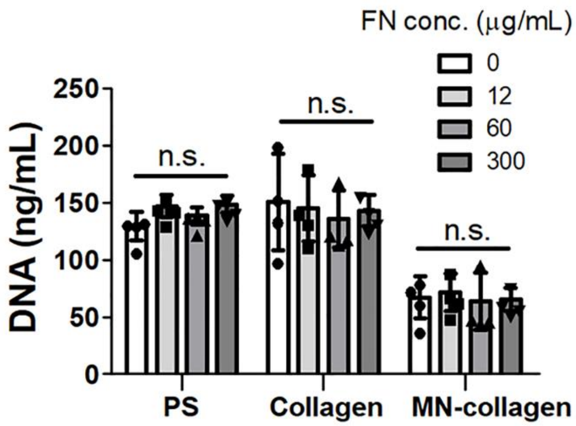


**Supplementary Figure 6:** MDA-MB231 cell growth on the different substrates for 7 d. Substrates were incubated with various concentrations of fibronectin (FN) before cell seeding ($n$ = 4).

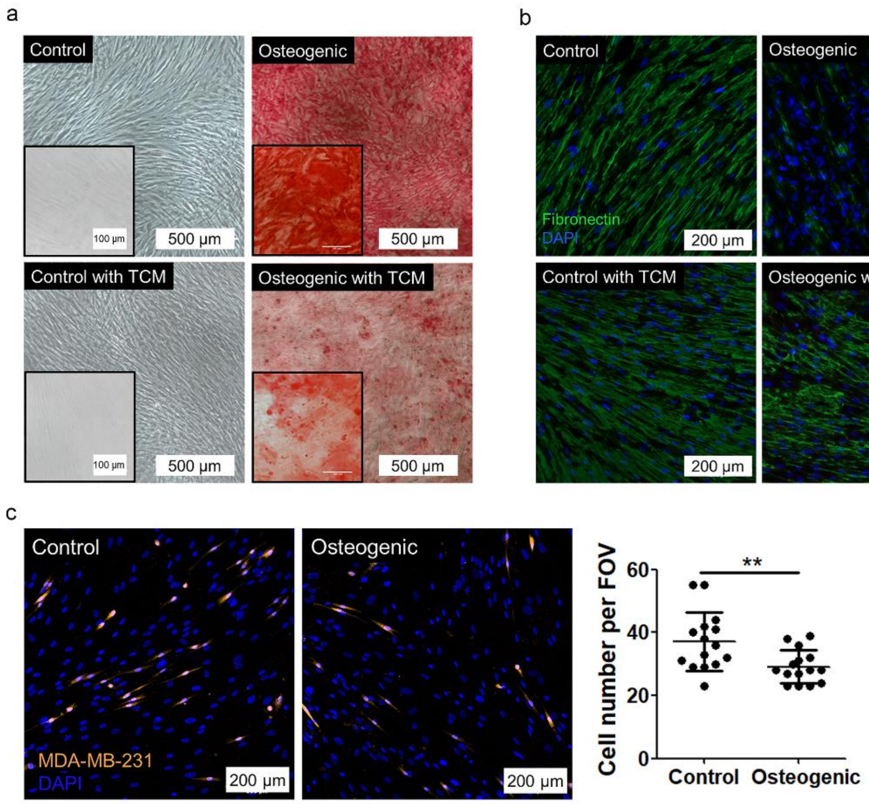


**Supplementary Figure 7: a,** Representative bright field micrographs of Alizarin Red S stained hMSCs cultured with control or osteogenic media and in the presence or absence of tumor-conditioned media (TCM). Scale bar = 500 μm. **b,** Representative confocal micrographs of hMSCs stained for fibronectin. Scale bar = 200 μm. **c,** Representative confocal micrographs and corresponding quantification of co-cultured osteogenically differentiated hMSCs and labeled MDA-MB-231 cells ($n$ = 3). Scale bar = 200 μm. **: $P < 0.01$ by non-parametric Mann-Whitney $t$-test.

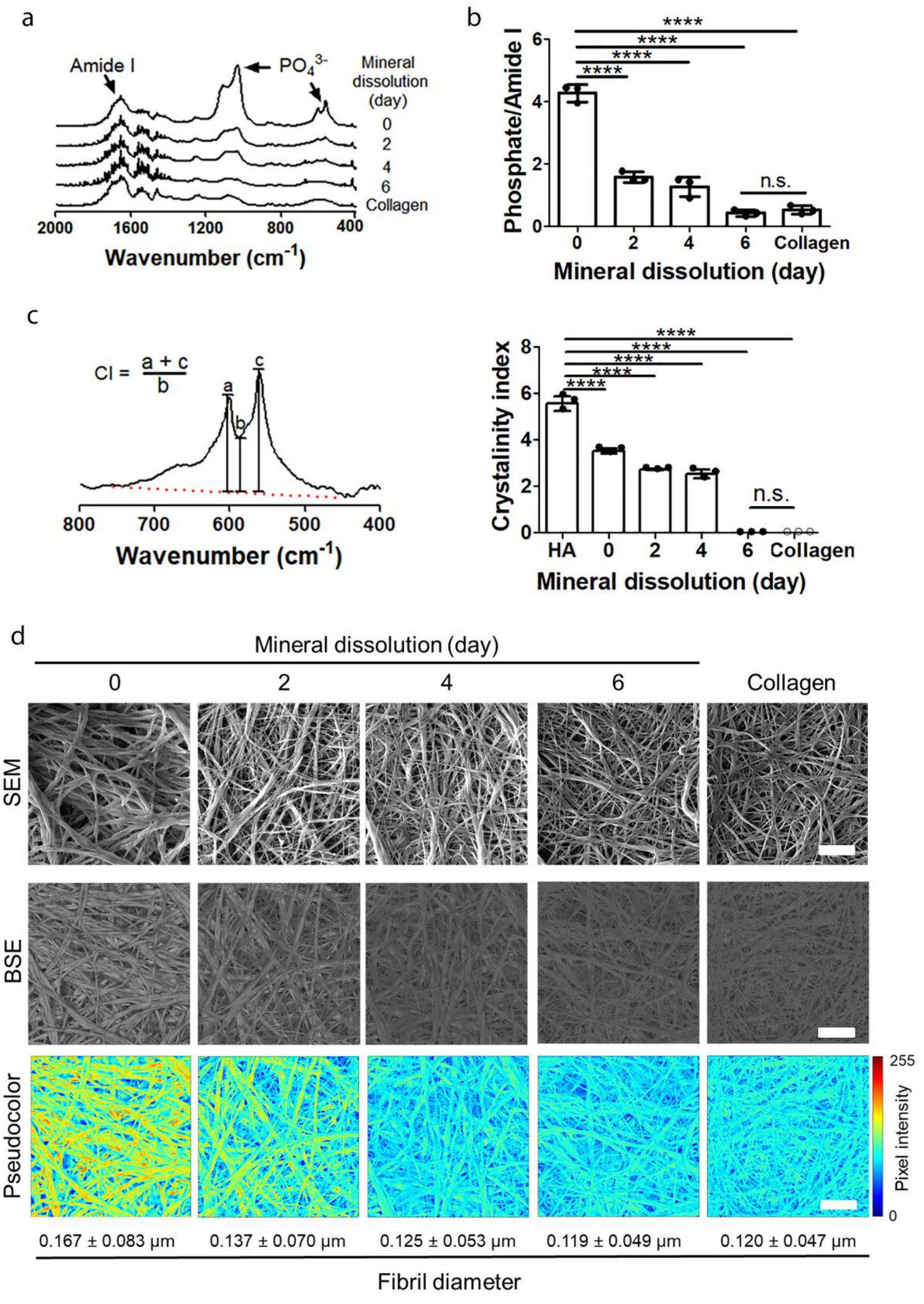
a
Amide I
$PO_4^{3-}$
Mineral dissolution (day)
0
2
4
6
Collagen
2000 1600 1200 800 400
Wavenumber ($cm^{-1}$)
b
Phosphate/Amide I
****
n.s.
0 2 4 6 Collagen
Mineral dissolution (day)
c
CI = (a + c)/b
a b c
800 700 600 500 400
Wavenumber ($cm^{-1}$)
Crystalinity index
HA 0 2 4 6 Collagen
Mineral dissolution (day)
d
Mineral dissolution (day)
0 2 4 6 Collagen
SEM
BSE
Pseudocolor
255
Pixel intensity
0
0.167 ± 0.083 μm
0.137 ± 0.070 μm
0.125 ± 0.053 μm
0.119 ± 0.049 μm
0.120 ± 0.047 μm
Fibril diameter

**Supplementary Figure 8: a,** FT-IR spectra of mineralized collagen following hydroxyapatite dissolution for up to 6 days. Non-mineralized collagen served as control. **b, c,** FT-IR analysis of mineral to matrix ratio (b) and crystallinity index (CI) of matrices after different periods of mineral dissolution (c) ($n$ = 3). CI was calculated by drawing a baseline between 450 $cm^{-1}$ and 750 $cm^{-1}$ (red dashed line) and measuring the heights of phosphate peaks at 567 and 603 $cm^{-1}$ and the height of the lowest point between them relative to the baseline. ****: $P < 0.0001$ by one-way ANOVA post-hoc Dunnett's multiple comparison test. **d,** Representative SEM and BSE images visualizing differences in mineral content and collagen fibril diameter after different periods of mineral dissolution. Pseudo-color was generated from BSE images. Fibril diameter was measured from SEM images. Scale bar = 2 µm.

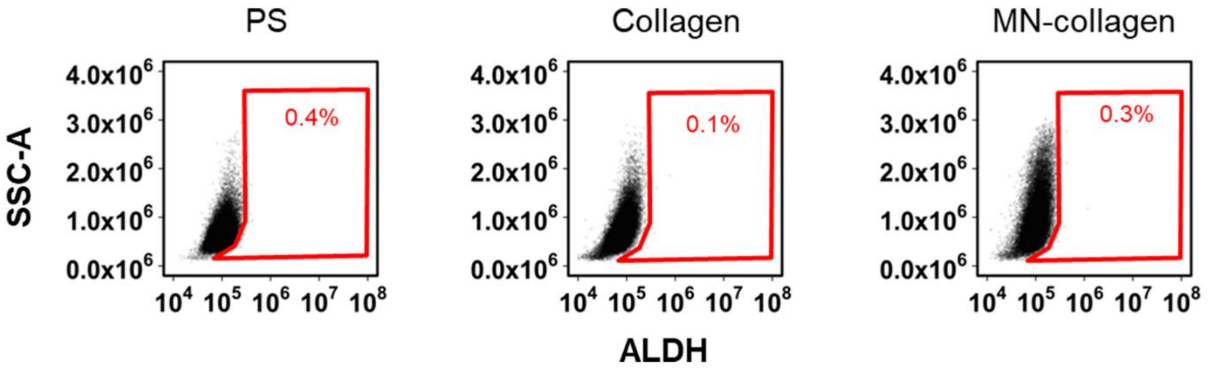


**Supplementary Figure 9**: Flow cytometry analysis of aldehyde dehydrogenase (ALDH) activity in bone metastatic subclone BoM-1833.

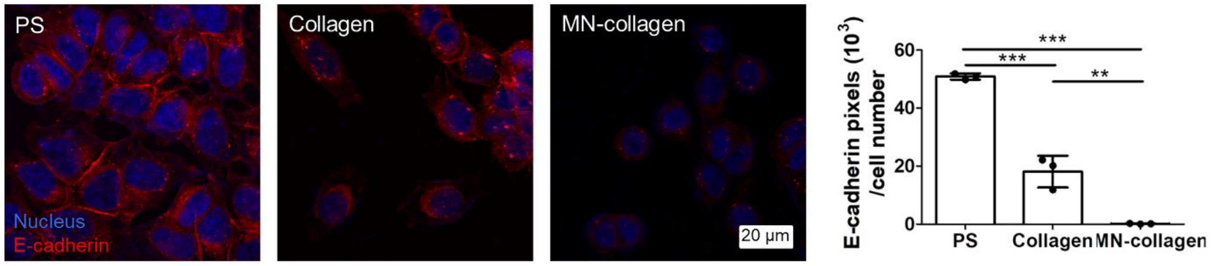


**Supplementary Figure 10:** Representative confocal micrographs and corresponding quantification of E-cadherin after 7 d of culture of MCF7 cells on the different substrates ($n$ = 3). Scale bar = 20 µm. **: $P < 0.01$, ***: $P < 0.001$ by one-way ANOVA followed by Tukey's post test for multiple group comparisons.

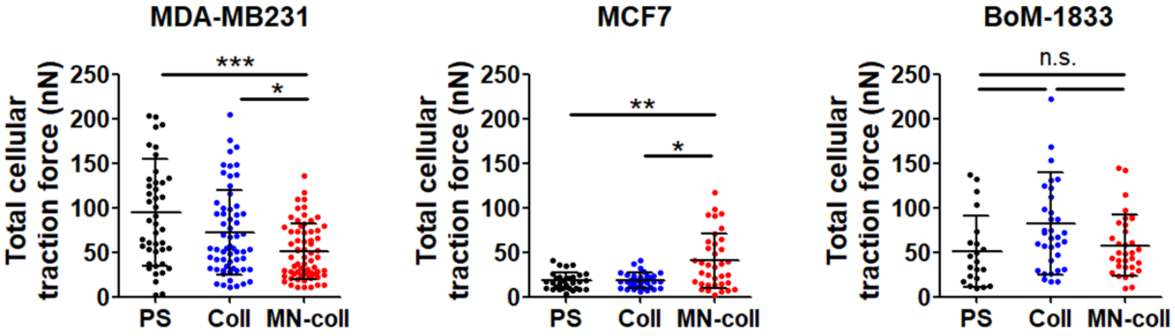


**Supplementary Figure 11:** Total traction forces of different breast cancer cell lines precultured on tissue culture polystyrene (PS), collagen, and mineralized collagen. Cells were precultured on the different substrates for 7 days prior to reseeding on PA gels for traction force microscopy measurements. *: $P < 0.05$, **: $P < 0.01$, ***: $P < 0.001$ by Kruskal-Wallis test with Dunn's multiple comparison correction.

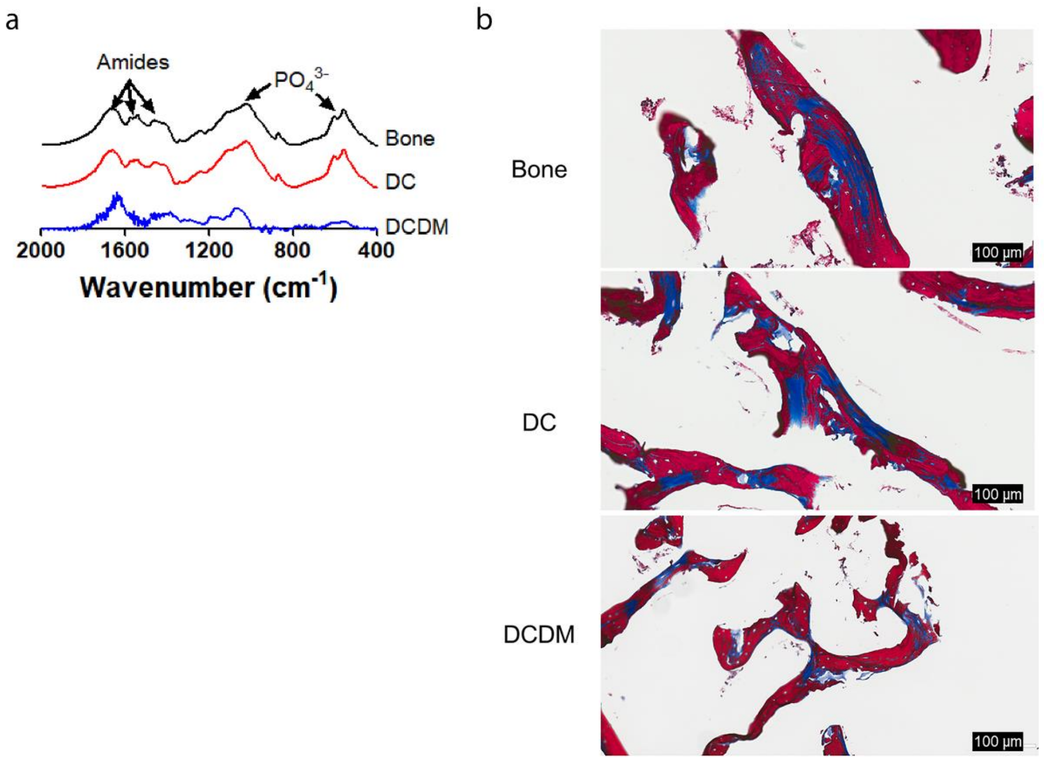


**Supplementary Figure 12: a, b,** FT-IR spectra (a) and Masson's trichrome staining (b) of DC and DCDM scaffolds and native bone. Scale bar = 100 µm. Masson's trichrome staining was performed after the removal of mineral for sectioning of scaffolds. Collagen (blue) and Bone (red).

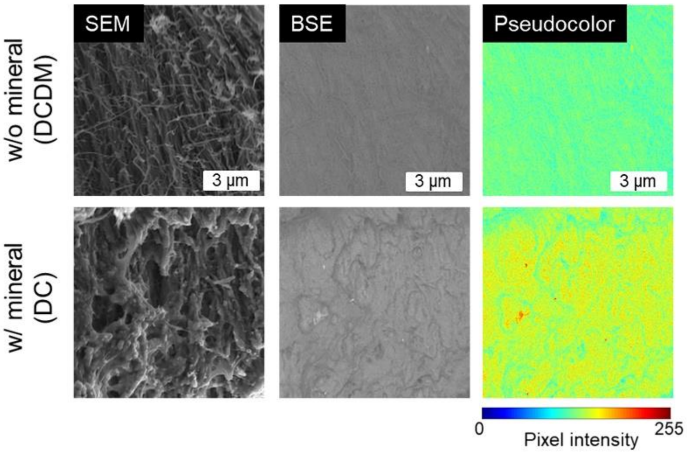


**Supplementary Figure 13:** SEM and BSE images of scaffolds after DCDM treatment. Pseudocolor indicates mineral content as determined from BSE. Scale bar = 3 µm.

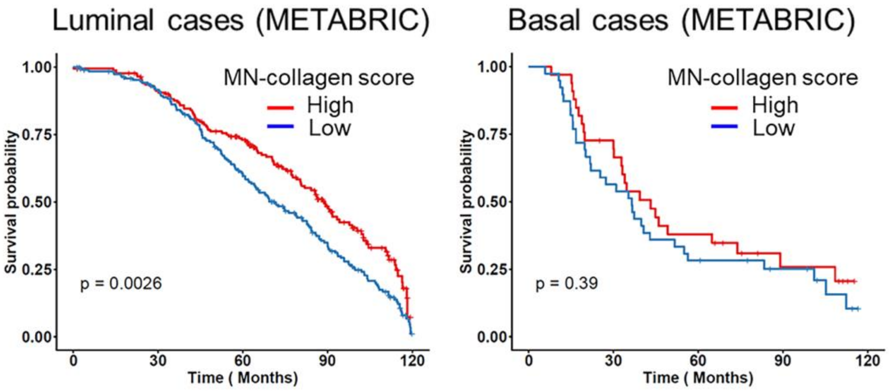


**Supplementary Figure 14:** Overall survival of breast cancer patients by stratification based on mineral-induced gene expression using different breast cancer subtypes in METABRIC database.

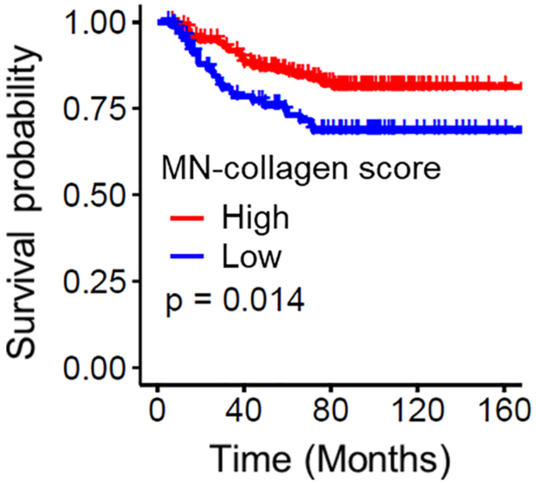


**Supplementary Figure 15:** Overall bone metastasis-free survival of breast cancer patients scoring high or low for expression of the mineral-induced gene signature (Table 2).

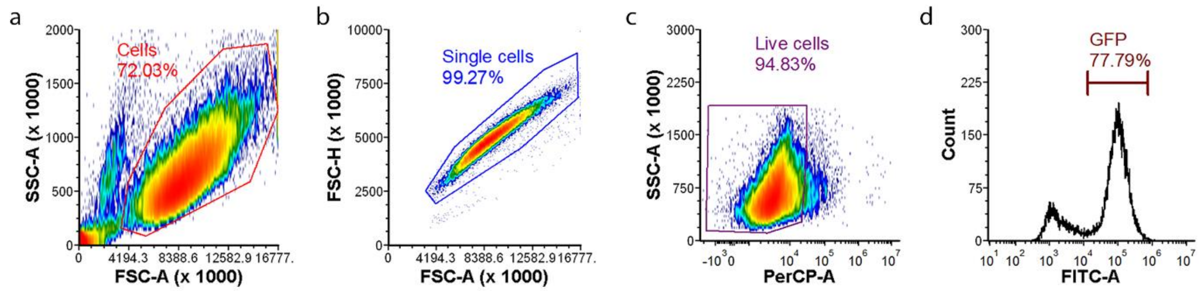


**Supplementary Figure 16: Flow cytometry gating strategy in this study.** Cells were gated by forward scatter area (FSC-A) and side scatter area (SSC-A) (a), then FSC-A and forward scatter height (FSC-H) signals were used for doublet exclusion to select single cells (b). Live cells were gated by propidium iodide signals (PerCP-A) and SSC-A (c). Cells populations expressing fluorescent signals (e.g. GFP from Nanog-GFP or ALDH from MDA-MB231) were analyzed (d).
